# Going with the flow: leveraging reef-scale hydrodynamics for upscaling larval-based restoration

**DOI:** 10.1101/2024.11.12.623286

**Authors:** Marine Gouezo, Clothilde Langlais, Jack Beardsley, George Roff, Peter Harrison, Damian P. Thomson, Christopher Doropoulos

## Abstract

Anthropogenic pressures are impacting coastal marine ecosystems, necessitating large-scale interventions to accelerate recovery. Propagule-based restoration holds the potential for restoring shallow coastal systems at hectare scales by harnessing natural dispersal. However, predicting propagule dispersal remains challenging due to the complex hydrodynamic nature of coastal marine ecosystems and the complex behaviours of marine propagules. To improve predictions of fine-scale larval dispersal patterns, we developed a 3D reef-scale (∼30 m resolution) dispersal model for Lizard Island, Australia, with the aim to predict the effect of island scale hydrodynamics on the distribution of coral spawn slicks and larvae. Using *in situ* field observations, and dispersal simulations, we assessed the model’s capability to (1) forecast hydrodynamic conditions, (2) predict coral spawn slick convergence zones for collection efforts, and (3) identify optimal locations and timeframes where high particle residence time may enhance local settlement following larval delivery to damaged reefs. Predictions of convergence zones in the upper water column aligned well with field observations of coral spawn slicks. At the reef benthos, the model captured variability in current speed and direction at ∼58% of studied locations. At other locations, the model did not resolve hydrodynamic conditions due to sheltering effects and associated hydrodynamic processes occurring at a scale below 50 m. At locations where the model performed well, propagules could remain within a one-hectare area around the delivery site for 5 to 15 hours depending on locations and the timing of larval release. These high retention conditions were infrequent but occurred at least once at 15 of the 25 studied sites. Observations of local currents *a posteriori* confirmed model predictions, showing periods of little water movement lasting from 6.5 to 15 hours. Overall, our study highlights fine-scale dispersal modelling as a key tool for scaling up larval-based reef restoration, while also acknowledging the need for better predictions of local conditions in complex, shallow environments. Applications of fine-scale modelling, coupled with local knowledge of reproductive timing and larval behavioural ecology, assist with the mass collection of propagules upon release and in identifying areas and times of optimal larval deployment to achieve the greatest impact.

## Introduction

Anthropogenic pressures increasingly impact coastal marine ecosystems (Halpern et al. 2007, 2008) at both global (ie. increasing sea-surface temperature) and local scales (ie. habitat destruction) (Hoegh-Guldberg 1999, Orth et al. 2006). Recent assessments have calculated damage or loss of large areas of marine habitats in the order of 10 to over 1 million hectares (Sudo et al. 2021, Asner et al. 2022). Many of these marine ecosystems are declining at alarming rates (Orth et al. 2006, Beck et al. 2011, Hughes et al. 2017), making it clear that protective management alone is no longer sufficient (Saunders et al. 2017) necessitating interventions that facilitate early recovery and restoration (Duarte et al. 2020, Randall et al. 2020). While new restoration techniques have been developed, their application is often limited to relatively small spatial scales across all marine coastal ecosystems, typically smaller than a few hectares (Bayraktarov et al. 2016). However, reviewed evidence of ‘bright spots’ demonstrates that restoration interventions can be effective at spatial scales exceeding 1,000 ha (Saunders et al. 2020).

A promising technique to upscale coastal marine restoration involves the mass deployment of propagules such as seagrass seeds, kelp spores, or benthic invertebrate larvae into damaged habitats to initiate early recovery (Vanderklift et al. 2020). The abundant propagule output of these organisms following reproductive events (e.g. Oliver and Willis 1987) offers unique opportunities to mass-collect propagules that are genetically diverse with low impact on adult populations (Broadhurst et al. 2008). Propagule collection and deployment are likely to be one of the approaches that have the potential to be applied at ecologically relevant medium to larger scales (≥1ha) (Rinkevich 1995, Doropoulos et al. 2019a, Banaszak et al. 2023). For example, eelgrass seeds were deployed across 125 ha in Virginia in the late 1990s and expanded to >1700 ha just 11 years later (Orth et al. 2012). For coral reefs, however, while the mass deployment of coral larvae onto reefs has been conceptualized (Heyward et al. 2002, Doropoulos and Babcock 2018, Doropoulos et al. 2019b, 2019a), and shown to be effective at restoring breeding populations within 2-3 years at small scales (dela Cruz and Harrison 2017, Harrison et al. 2021), empirical demonstration of restoration success at hectare scales is currently lacking.

The principal challenge in upscaling larval-based restoration of benthic marine invertebrates lies in identifying (1) where to collect gametes and developing embryos *in situ* following reproductive events and (2) when and where to release competent larvae to significantly increase local larval settlement and recruitment on target reef areas. This requires understanding the physical properties of propagule dispersal at very fine spatiotemporal scales (i.e. minutes to hours and ≤ 50-100 m) in conjunction with propagules’ biology and behaviour occurring during early development in fish and invertebrate larvae (Bode et al. 2024). Recent research has shown that using dyes or colouring the coral propagules (Doropoulos and Roff 2022) can assist to better predict and observe the dispersal of invertebrate propagules at small spatial scales (i.e., hundreds of meters). Additionally, employing fine-scale dispersal modelling during interventions can help predict local circulation conditions at the study location (e.g. Bruyère et al. 2023). When coupled with an understanding of larval biology and ecology (Oliver and Willis 1987, Connolly and Baird 2010), dispersal modelling can guide where and when to collect propagules and release propagules once they are competent to transition as larvae in the water column to sessile organisms on the benthos.

Biophysical modelling plays a crucial role in benthic ecology, with applications ranging from global and regional studies that explore connections among ecosystems (e.g. Wood et al. 2016, Hock et al. 2017, Thompson et al. 2018), conservation and aquaculture practices (e.g. Boschetti et al. 2020, Bruyère et al. 2023), down to small-scale analysis of water movement in lagoons or bays (e.g. Wolanski and King 1990, Golbuu et al. 2016). Given the utilisation of biophysical models in operationalising real-world decision-making (Weeks 2017, Bode et al. 2019), a detailed assessment of circulation and validation of dispersal model predictions is required. Modelling accurate circulation and propagule dispersal at sub-mesoscales is challenging (Griffies et al. 2009, Fox-Kemper et al. 2019). Firstly, complexity arises from the multitude of factors influencing local hydrodynamics in shallow benthic habitats, including wind-driven currents, seafloor bathymetry, tidal forces, coastal inflow, swell and waves, and localized weather patterns (Wolanski et al., 2024). These factors operate at spatiotemporal scales often finer than observed hydrodynamic conditions used in hydrodynamic models (Largier 2003, Nickols et al. 2012, Morgan et al. 2018, Swearer et al. 2019, Wolanski et al. 2024). For example, coral reefs may experience significant differences in current speeds among different habitats within a reef system at a given hour, varying up to 16-fold (Johansen 2014, Philipps and Bellwood 2024). Secondly, marine propagules, which are small (typically less than 1 mm for broadcast coral spawners and less than 4 mm for seagrass species but larger for mangroves), exhibit varying buoyancy and behaviour throughout their dispersal phase These characteristics interact with hydrodynamics and thus need to be integrated into. Lagrangian particle tracking models (Swearer et al. 2019). For example, buoyant propagules floating at the surface are generally pushed directly downwind by both currents and wind drag (Van der Mheen et al. 2020). However, this wind drag effect is absent for propagules that remain sub-surface (Van der Mheen et al. 2020). Validating modelled hydrodynamics and predicted dispersal patterns of propagules is essential for understanding model inaccuracies, assessing confidence levels in predictions, and identifying limitations for future improvements. However, such validations are rarely prioritized. When included, validation of modelled circulation and dispersal patterns has so far included *in-situ* measurements of current conditions to assess hydrodynamic models (Dumas et al. 2012, Bruyère et al. 2023), collecting field data on propagule supply and settlement to assess dispersal patterns compared with biophysical model predictions (Schlaefer et al. 2018, Gouezo et al. 2021, Doropoulos et al. 2022), or utilizing genetic parentage data to assess how populations are connected (Riginos et al. 2019, Bode et al. 2019).

Coral larval restoration involves collecting embryos, culturing larvae, and then delivering competent larvae to damaged reef areas (Heyward et al. 2002, dela Cruz and Harrison 2017, Doropoulos et al. 2019b, Harrison et al. 2021). Embryo collection from coral spawn slicks occurs mainly at night following the synchronous, mass release of eggs and sperm into the water column (Harrison et al. 1984). Coral spawn remains positively buoyant for the first 12 to 16 hours (Harrison et al. 1984, Oliver and Willis 1987, Wolanski et al. 1989). During this buoyant phase, dispersal is influenced by ocean currents (fronts, internal tides and waves), wind conditions, and coastline and reef topography (Wolanski et al. 1989, Pattiaratchi 1994, Smith et al. 2021). This transport leads to their accumulation in convergence zones, known as Langmuir circulation, and form visible slicks at the surface (Wolanski and Hamner 1988, Wolanski et al. 1989, Dethleff et al. 2009). Identifying the convergence zones with high propagule concentrations is important for efficiently collecting coral spawn or deciding

where to deploy passive spawn catcher systems (Harrison and dela Cruz 2022). Once coral gametes and developing embryos are collected, they are cultured in culture pools or tanks on vessels until larvae reach competency (e.g. Heyward et al. 2002, Doropoulos et al. 2019b). Once competent (and neutrally buoyant), larvae are dispersed onto damaged reefs to initiate early coral recovery. Currently, delivery is conducted under tents or plankton mesh sheets to constrain larval dispersal (Heyward et al. 2002, Edwards et al. 2015, dela Cruz and Harrison 2017, 2020, Harrison et al. 2021). This method is highly effective but limits the spatial scale of operations to <100 m^2^. By understanding and modelling local hydrodynamics and dispersal patterns, we can better select where and when to release coral larvae to maximize local retention, potentially scaling up coral reef restoration to hectare scales.

Here, we apply a reef-scale model (∼30 m resolution) to predict the transport and dispersal of coral spawn and coral larvae. The reef-scale hydrodynamic and dispersal findings can be used to inform decisions during larval restoration interventions such as selecting locations to collect coral spawn samples or deliver competent larvae. First, we assessed the predicted currents of the hydrodynamic model using *in situ* observations of current conditions, both near the surface and at depths, to highlight the strengths and weaknesses of our approach and the limitations of the hydrodynamical model (Objective 1). We then combined *in situ* current datasets, field observations, and particle dispersal simulations to assess the usability of biophysical models in reef restoration. For the collection phase, we modelled the dispersal of buoyant coral gametes following spawning events to predict the occurrence of convergence zones and relate predictions to field observations (Objective 2). For the delivery phase, we explored the occurrence and variability of local retention conditions on reefs using modelled residence time of particles within a hectare of a delivery location during the entire delivery timeframes to predict optimal times for larvae delivery on damaged reefs (Objective 3). We then contrast findings *a posteriori:* comparing modelled residence time with slack current conditions data collected using *in situ* tilt current meters during delivery timeframes at target deployment reefs during interventions (Objective 4). Lastly, we discuss the potential for utilising reef-scale hydrodynamics to upscale coral larval restoration, limits of the model, and key knowledge gaps for future research.

## Methods

### Study location and high-resolution dispersal model

The study was conducted at Lizard Island, Northern Great Barrier Reef (GBR), Qld, Australia (14.6645° S, 145.4651° E) (Fig. S1). Lizard Island and the surrounding reefs are situated in the shallow water (0-20 m depth) mid-shelf of the GBR, influenced by hydrodynamic conditions typical of mid-shelf islands, largely protected from open ocean swell (Fulton and Bellwood 2005, Madin et al. 2006). The reef habitats around Lizard Island include sheltered bays, lagoons, semi-exposed lagoonal habitats, channels, and exposed outer reef slopes (Roelfsema et al. 2014), making the location ideal to test current hydrodynamic and larval dispersal models.

To reach a resolution of ∼30-50 m around Lizard Island’s reef, two nested 3D hydrodynamic models were developed using the Environment Model Suite (EMS) (Baird et al. 2020): a doughnut shape domain nested inside the 1/10° global model OceanMAPS v4.0 (Australian Bureau of Meteorology (ABOM) 2017, Schiller et al. 2020), and a 50 m regular-grid domain nested inside the donut-shape domain. (Fig. S1). The high-resolution model domain covers approximately 8 by 8 km, and the modelled current can be used to drive the particle tracking model CONNIE (Condie et al. 2012). Atmospheric forcing is provided by the ACCESS-G atmospheric model (Puri et al. 2013).

CONNIE allows the release of a set number of virtual particles within chosen polygons at a chosen depth. Particles are released randomly within polygons during the hour of the chosen release. From their release, each particle is then tracked through space. The outputs of CONNIE dispersal simulations are in JSON format showing the locations of each particle through space (latitude, longitude, depth) at each time step during the chosen dispersal timeframe. For this study, particles are bound to their release depth. The modelled ocean currents are used to advect the particles, accounting for wind-driven ocean circulation and tidal circulation. For particles released near the surface, CONNIE has the option to add a surface wind drag forcing set as a % of wind conditions. The hydrodynamic model and particle tracking model CONNIE ran at a 12-minute time step in near real-time, with the option of a 3-6 day forecast to apply the workflow planning to periods ahead of real-time coral larval restoration interventions to support decision making. This modelling approach was designed to enhance the precision and efficiency of coral larval restoration, ensuring targeted and effective collection of coral spawn slick samples for mass larval culturing and deployment of competent coral larvae to maximize the success of reef restoration initiatives.

### In-situ hydrodynamic data collection

Throughout the study, particularly during coral larval intervention timeframes, *in situ* hydrodynamic data were collected around Lizard Island’s reefs with the two main goals: (1) to assess localized current variability and validate model predictions in the context of coral larval restoration, and (2) to identify potential model weaknesses and limitations occurring at such fine resolutions for future improvements.

Hydrodynamic conditions near the surface (∼0.5 to 1 m below the surface) were captured using surface drifters (Pacific Gyre Microstar drifters) through n=19 runs within the model domain during spawning events in November 2021 and December 2022. The drifters operate at 1 meter depth and were set to record the near surface current drift by emitting their GPS locations at 5-to-10-minute intervals. This *in situ* hydrodynamic dataset was used to compare drifters tracks to the modelled drift of particles at the scale of the island (see next section: *Model validation approach*).

Hydrodynamic conditions below the surface (>2 m depth) were collected using stationary tilt current meters (LOWELL TCM-1 Tilt Current Meter) to quantify current conditions in the water column layer of ∼25 to 100 cm above the benthos and within a 1 m radius surrounding where they are attached to the reef. Tilt current meters were set to record current speed and direction at 1-second intervals at a total of 28 reef locations with depth ranging from 2 to 9.5 m. Data were recorded over a combined total of 6 weeks spread over 2 years, with a focus on conditions during larval deployment interventions conditions (i.e. November-December 2021 and 2022). Time series at each station were then averaged at longer time scales for analyses. This *in situ* hydrodynamic data set was used to assess the ability of the model to accurately predict conditions at a reef site scale, often located at shallow depths and close to the coastline, to best inform local conditions during larval delivery timeframes (see next section: *Model validation approach*).

Given that the hydrodynamic model grid cells have a minimum resolution of ∼30-50 m, the model bathymetry does not represent the complexity of the reef structure nor the resulting small-scale hydrodynamic patterns. For this reason, in this study, we attached tilt current meters at the top of reef boulders or reef, referred to as “reef top” so that the loggers captured near-bottom flow that the model can represent.

To gain a better understanding of the variability in current speed and direction occurring within a grid cell and may be influenced from reef structural complexity, we measured the current at 11 locations within a model grid cell over a 24-hour cycle. This was achieved by deploying 11 tilt current meters in a 30 x 30 m square on the reef, at two sites: a structurally complex reef slope with boulders, overhangs and >40% coral cover, and a flat rubble bed with <10% coral cover and no boulders. At the reef slope, half of the current meters were deployed on top of the reef structure (ie. reef tops) and half in locations in between boulders, referred to as “reef grooves”, where structural surroundings would either block the dominant flow and/or likely create small eddies and turbulences. At each of these two locations, we visualized the variability in current speed (averaged at 1 min interval) and direction (distribution frequency of current headings) among the 11 tilt current meters. The impact of complexity (categorized into two levels: reef tops and grooves) on 1 min averaged current speed was assessed using a linear model. The current speed response variable was fourth root-transformed to conform to the normality of model residuals.

### Model validation approach and assessment of localized current variability

To validate the modelled hydrodynamic conditions near the surface to provide information on coral spawn slick convergence zones, tracks from 1,000 particles were released within a 50 m radius polygon around the starting location of each drifter run (n=19) and compared with drifter tracks. Since the drifters used in the study captured the drift within the top 1-m layer of the water column from the surface, particle dispersal scenarios were conducted at the four modelled depths of 0.25, 0.75, 1.3, and 1.95 m, and no wind-drag forcing was applied. A spatio-temporal analysis between drifter and mean particle tracks was conducted by calculating the error in distance, direction, and speed between the two tracks every hour. Drifter and particle tracks were visualized in QGIS (v3.34.1) (QGIS Development Team 2023), and spatiotemporal analysis between tracks was performed in R using the Rgdal, spacetime, and sp packages (Pebesma 2012, Pebesma et al. 2012, Bivand et al. 2015), following a similar workflow as in Santos et al. (2016). The influence of depth on the distance error and direction error between particles and drifter tracks was tested using a linear model. Model residual normality and variance of homogeneity were checked to determine whether model assumptions were met.

To assess localized current variability between *in situ* and modelled current and validate the modelled hydrodynamic conditions below the surface (>2 m depth) for delivery dispersal scenarios, time series of current data predicted by the model were compared to current data collected by stationary tilt current meters. Modelled data were extracted from the grid cell nearest the location of the tilt current meter. In cases where the grid cell nearest the location of the tilt current meter was significantly shallower or deeper than the tilt current meter (i.e. 2 m) (i.e., due to small bathymetric errors caused by resolution), the nearest grid cell with a better bathymetric match was chosen. Time series of both modelled and *in situ* u and v velocity vectors were then averaged hourly and compared using linear regression analyses of modelled u and modelled v against observed u and observed v, respectively. Scatter plots were generated, and R^2^ and root mean square error (RMSE) were calculated for each reef site and plotted to visualize potential model domain area with better accuracy (Fig. S2-8).

After this evaluation, reef locations were categorized into four groups: (1) “very good fit,” indicating strong agreement for both velocity vector components, speed, and direction (R^2^ >0.4, low RMSE value), reflecting an accurate representation of wind-driven circulation, tides, and local effects; (2) “good fit,” demonstrating satisfactory agreement or acceptability for one of the velocity vectors, current speed, and direction. While the model may miss some local effects or experience lag issues with tides, these limitations are well-understood, and expert knowledge can compensate for them, allowing the model to inform decision-making. (3) “poor fit” shares characteristics with the “good fit” category, but the limitations cannot be overcome by expert knowledge, rendering the model unsuitable for decision-making. (4) “no fit” category: the model fails to represent observed local processes and dynamics. This selection criterion aimed to assess where the model simulated precise hydrodynamic conditions that are known to be highly variable near the benthos while allowing room for expert interpretation of predicted patterns.

### Collection phase: predicting coral spawn slicks

Based on the findings of the validation analysis, coral spawn slicks were simulated at 0.25 m depth only, with and without wind drag drift forcing. Considering that surface wind forcing on floating coral gametes is poorly understood (Oliver and Willis 1987, Pattiaratchi 1994, Van der Mheen et al. 2020), and acknowledging the likelihood that Lizard Island creates a wind shadow in the lee of the island, we tested two surface wind drag drift forcing scenarios on particles near the surface: 0 and 3% of the wind speed, the latter commonly used to simulate oil spills or dispersion of seagrass seeds at the surface (Erftemeijer et al. 2008, Le Hénaff et al. 2012). The modelling exercise helped visualize the potential effect that the wind drag drift parameter can have at a reef system-scale at the beginning of the dispersal phase after spawning (i.e., 0-16 hours) when particles are extremely buoyant (Oliver and Willis 1987).

To enhance the accuracy of modelling coral spawn slicks, we accounted for spatial variability in production of coral spawn and considered the relative influence of reef area and live coral coverage on particle density. Particles were dispersed within polygons containing live corals with ≥10% cover, using a combination of habitat maps, the Allen Atlas, AIMS LTMP data (Roelfsema et al. 2014, Allen Coral Atlas 2022, AIMS 2023), and our own *in situ* observations. The relative relationship between larval density and coral coverage was calculated following Roff (2023) allowing estimates of the relative number of particles within each polygon based on its area and live coral coverage (Fig. S9). Subsequently, polygons were divided into approximate portions of predicted densities of ∼5,000 particles to simplify the modelling exercise so that a set number of particles of 5,000 were dispersed within each polygon (Fig. S9). This resulted in 90 polygons of various areas based on their cover (Fig. S8), each with 5,000 particles released, totalling close to half a million particles dispersed per spawning event.

Particle dispersal was simulated during conditions from three observed spawning nights in 2021 and 2022, specifically 21-23/11/2021 and 09-11/12/2022. Particles were dispersed near the surface at the modelled depth of 0.25 m from 19:00 until 21:00 and were tracked for 16 hours, including with and without a surface wind forcing of 3%. The .tiff output of each dispersal simulation was summed in QGIS for each scenario to represent the overall density of particles within the model domain during the 16-hour dispersal timeframe. This allowed us to identify zones of slick convergence, informing where collection efforts should be focused.

### Larval delivery phase: predicting the residency of particles over damaged reefs

The dispersal of passive particles at 25 delivery locations during delivery time periods was conducted to: 1) quantify the residency of particles over the reef they were delivered onto and within a hectare (in all directions) from the delivery source; (2) investigate how dynamic particle residency is depending on when during the day they are delivered; and, (3) identify locations that, due to their surrounding hydrodynamic features, may have properties that enhance retaining or aggregating particles within one hectare from the delivery source. Such locations are ideal for releases of larvae, as the longer particles remain over the reef, the higher the likelihood they will settle ‘en masse’ and promote early coral recovery. These simulations were conducted to quantify the residency of particles over reefs over 24 hours, directly related to hydrodynamic forces and physical reef features. Thus, particles were considered neutrally buoyant for this exercise, and the prediction of the likelihood of settlement based on multiple additional factors (larval biology and behaviour, receiving environment, presence/absence of reef, etc.) is the focus of another study.

Particle dispersal was simulated within 25 polygons (25m radius) within the model domain that showed acceptable validation findings (Objective 1). In each polygon, 5,000 particles were released at a depth of 3.6 m every hour from 6:00 until 16:00 over five days in both 2021 and 2022. As the timeframe for coral larval delivery typically ranged from 4 to 8 days following a spawning event (i.e., approximately 6-10 days following the full moon), these timeframes in 2021 and 2022 were selected to be from 27-30/11/2021 and 13-17/12/2022. The depth of released particles was based on a typical operational depth during larval deployment on damaged reefs, avoiding the surface layer (0 - 2 m deep) where the wind can exert a strong influence. Passive particles were tracked for 24 hours, as beyond that point, many were likely to hit the model boundaries, where they are considered ‘lost’. Conducting 11 simulations per site over 5 days in both 2021 and 2022 resulted in a total 2,750 individual particle simulations. Particles dispersal outputs (.json files) at 12 min time steps for 24 hr were batch post-processed in R using jsonlite, sf, and tidyverse R packages (Ooms 2014, Wickham and Wickham 2017, Pebesma 2018) to extract the number of particles that remained within approximately one hectare surrounding the delivery source (a ∼200 m radius buffer from the centre of the delivery source), ensuring that each particle was always mobile. Particles that exhibited stationary behaviour lasting >1h due to model artefacts were removed (Supplementary Information Appendix 1)

For each scenario and site, the sum of particles was calculated at every time step (i.e., 12 min) and then every hour, providing data on particle concentration through time at each site. Particle residency time (PRT) in hours within one hectare of the delivery source was calculated using the e-folding time, which is the time it takes for the initial concentration of particles (here, between 5,000 and 3,500) to drop to 1/e of its initial value (1,800 to 1288) within the defined area (Bolin and Rodhe 1972). This method is commonly used as a metric for particle retention within a defined location (Couto et al. 2017, Gouezo et al. 2021, Hudson et al. 2021).

PRT was compared among sites and through time using a negative binomial generalized linear mixed effects model with delivery site, delivery time, and their interaction as fixed factors, with delivery days included as a random effect in the model to account for the nested structure of the dataset using the glmmTMB package (Magnusson et al. 2017). Dispersion and model residuals were checked and validated using the package DHARMa. All statistical analyses were conducted in R version 4.3.2 (R Development Core Team 2023).

### *Larval delivery phase:* a posteriori in situ *current speed conditions*

To investigate current conditions on targeted reefs *a posteriori* of larval deployment interventions to meet Objective (4), 17 of the 40 available *in situ* current time series were selected. These time series, averaged at 1 min intervals, represent the current conditions at 17 reefs during larval delivery in 2021 and 2022. They encompass six habitats at depths ranging from 2 to 5 m.

First, differences in current speed among the studied sites were investigated using a generalized linear model with a gamma distribution to test for overall spatial differences among sites. Simulation-based model residuals were checked using the DHARMa package (Hartig 2017). Second, within each current speed time series, timeframes of slack current were selected using the changepoint package (Killick and Eckley 2014). Current speed trends were plotted, and timeframes of significant changes in current speed were identified for each site separately, adjusting the penalty value to modulate the sensitivity of changepoint selection (Fig. S10). Once timeframes of changes in current speed were detected, their start and end times, duration, average, and standard deviation were extracted. Third, timeframes of slack currents were selected, defined in this study as an average current speed usually <0.022m s^-1^. The frequency of the duration of slack current conditions was visualized to determine the most common duration. The count of slack current conditions at each site, along with their average duration and range of duration, was plotted to explore instances where larval retention at the reef would be maximized and dispersal minimized to promote local larval settlement. Lastly, the correlation between slack current conditions and the timing of high and low tides was examined to underscore the limited predictive capacity of tide timing in determining current strength. Timeframes of slack current and the timing of high and low tides were plotted for each site during each study period.

## Results

### Validation of the physical component of the model near the surface

Drifters runs confirmed a close match between modelled and in-situ current direction with 16 of the 19 drifter runs showing an acceptable match in drift direction when compared to particle dispersal tracks (Fig. 1.a.b.). The speed of drift between particles compared to the drifters was generally slower and less accurate than the direction, with a median error in speed of −0.036 m.s^-1^, in distance of 323 m and in direction of 42 degrees. The effect of particle depth (0.25, 0.75, 1.3, and 1.95 m) did not significantly influence the direction or speed of particle drifts (LM, P > 0.05).

**Figure 1:**
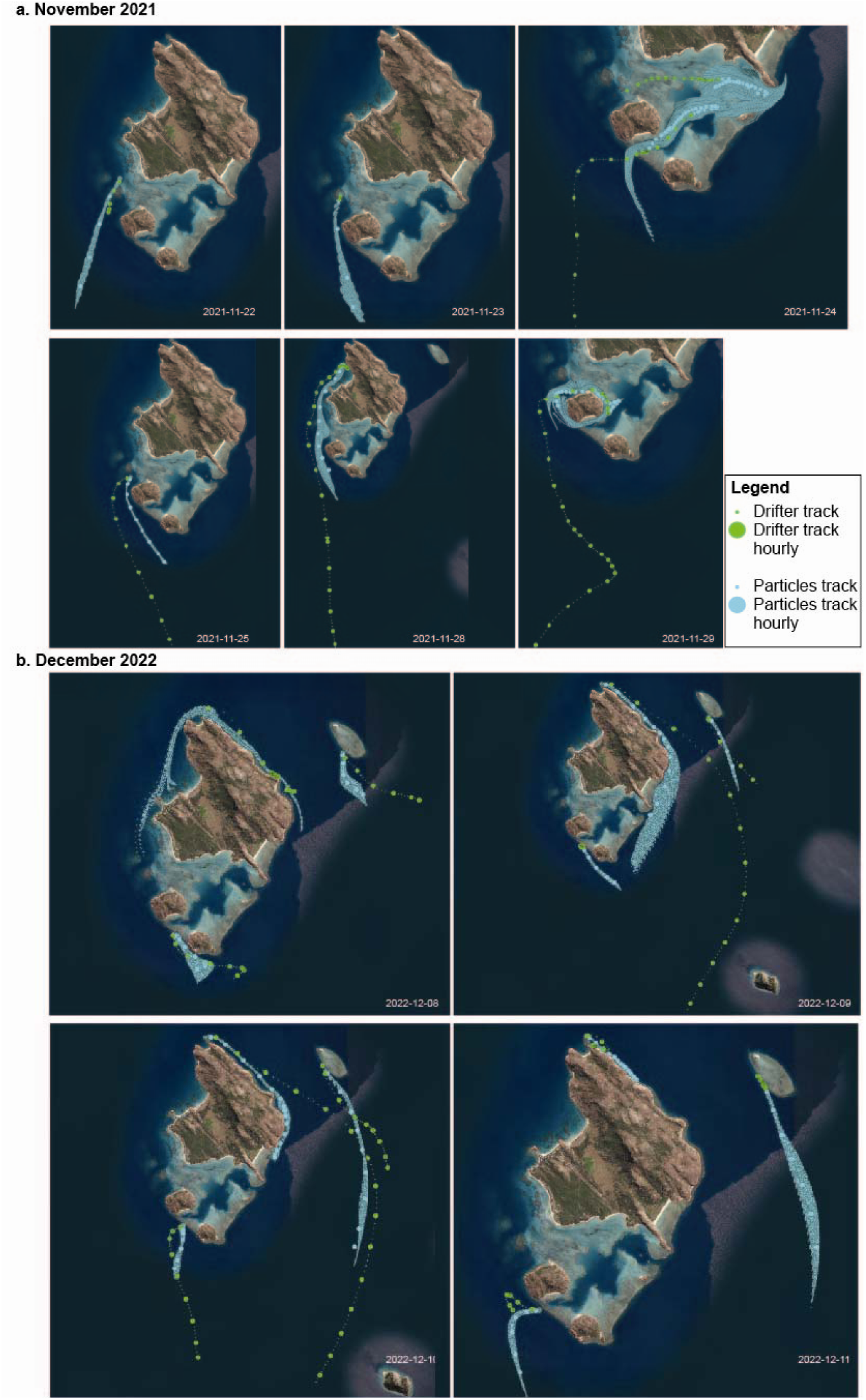
Maps showing tracks from surface drifters (green) and particles tracks from dispersal modelling (blue) during coral spawning season in 2021 (a) and 2022 (b)

### Predicting coral spawn slick dispersal near the surface for collection efforts

Dispersal of coral spawn slicks during spawning nights in both 2021 and 2022 exhibited considerable differences when including or excluding the surface wind drag forcing of 3% on particles (Fig. 2). Convergence zones with high particle concentrations, commonly observed in the field on key spawning nights and early mornings, were more pronounced when surface wind was excluded. In contrast, when wind was included, the plumes were much more diffuse and dispersed in a more westward direction, as opposed to the southward direction when surface wind drag was excluded.

**Figure 2:**
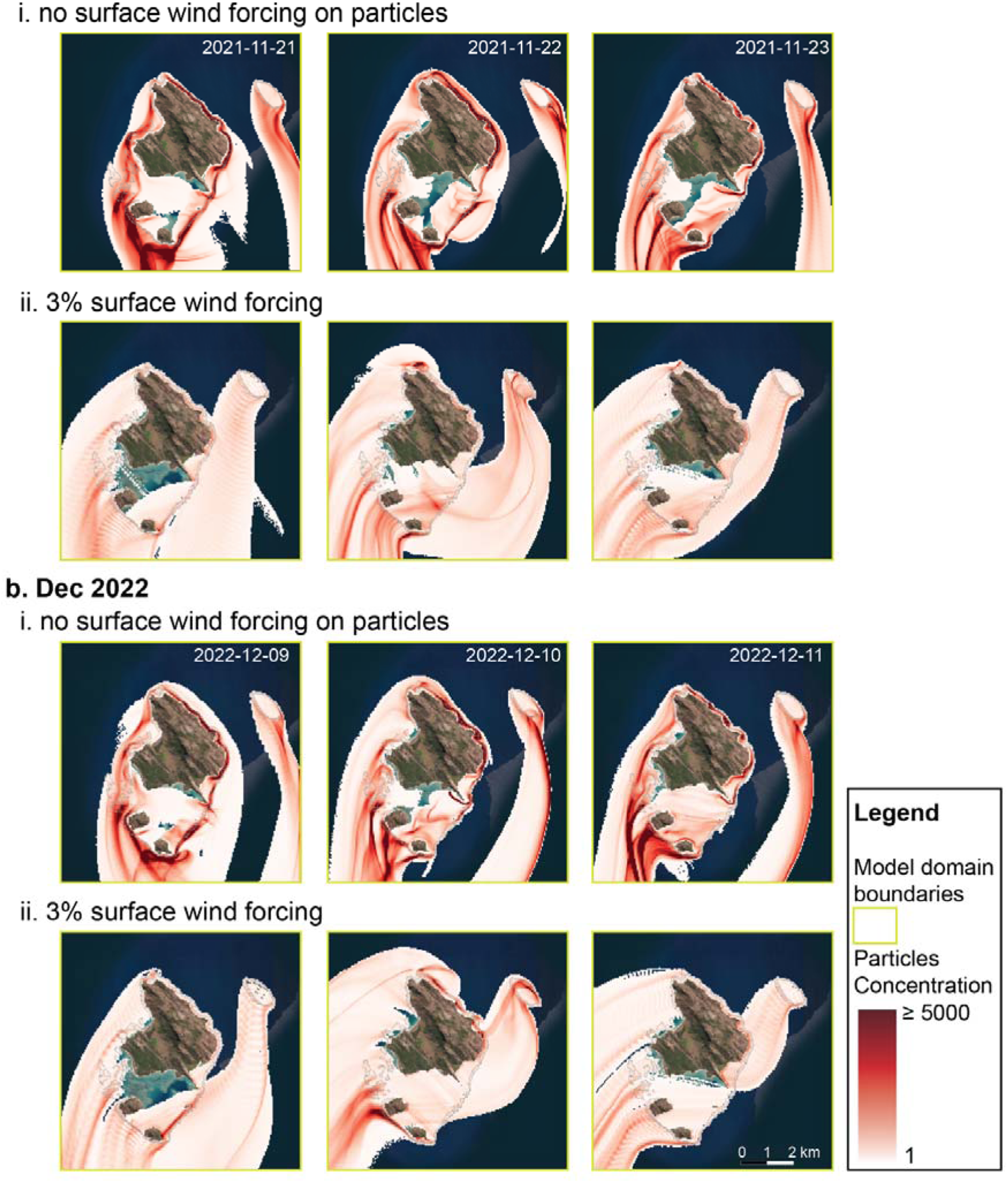
Coral spawn slicks dispersal showing the concentration of particles around Lizard Is. reefs over 16 hours following a spawning event in b 2021 (a) and 2022 (b) with (ii) and without (i) the surface wind forcing on particles dispersed.

Throughout the six spawning nights, particles consistently aggregated in ‘convergence zones’ and were primarily located in the region south of Palfrey Island reef system. This observation correlated with aerial observations of slicks and the collection of coral spawn using passive spawn catchers in December 2022 (Fig. 3).

**Figure 3:**
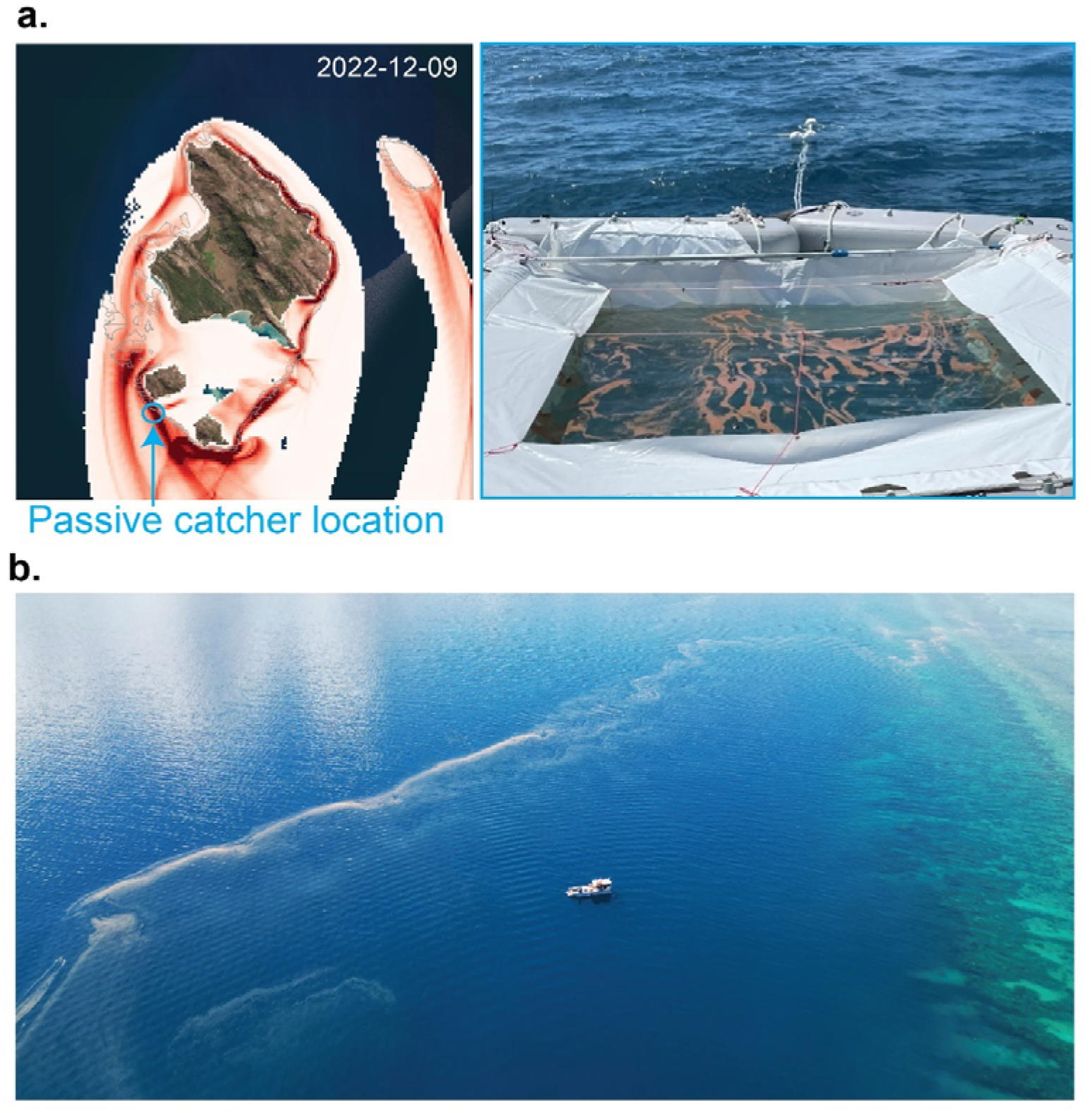
Early morning photograph of collected spawn in a passive spawn catcher (Harrison and dela Cruz 2022) placed along the outer reef slope between the two Palfrey islands (a), aerial photograph of coral spawn slicks along a fore reef observed in the mornings following major coral spawning events (b).

When surface wind drag was excluded, larvae plumes were less diffuse and overall, more concentrated near reefs. For instance, along the flanks of the north-eastern and eastern reefs, particles transported by the dominant southward current interacted with the presence of land and reef, which led to plumes being ‘stickier’ and particles retained near these reefs (Fig. 2). In contrast, in the open ocean south of McGillivray reef, stickiness was less apparent, and the diffusion of particles was much higher, irrespective of the effects of surface wind forcing (Fig. 2). This finding aligns with early morning observations of coral spawn slicks often observed to form along fore reefs (Fig. 3)

### Localised assessment of modelled and in situ current variability below 2m depth

#### Sub-scale spatial variability within a model grid cell

Current speeds recorded by tiltmeters deployed within a grid cell (30 x 30 m) were variable (Fig. S11). This variability was accentuated at the structurally complex reef, with overall significantly higher flow rate when the tiltmeters were positioned on top of the reef structure as opposed to down in the “grooves” (i.e., between two reef tops) (LM, P<0.001). The direction of current was also more variable in the grooves than on the reef top, implying the occurrence of small-scale turbulences. The variability in current speed and direction among tilt current meters was less evident at the rubble bed locations, which was close to flat.

#### Assessment of variability between modelled and *in situ* current data at reef site scale

Among the 40-time series of current data analysed (Table 1, Fig. 4, Fig. S2-8, Table S1), eight showed a very good fit, 12 exhibited an acceptable fit with limitations that could be addressed with expert knowledge, three showed a partially acceptable fit, which was contingent on wind conditions, while three had a poor fit that expert knowledge could not rectify. Additionally, 14 time series showed no fit, indicating that the present model could not resolve hydrodynamic processes occurring at the reef site.

**Figure 4:**
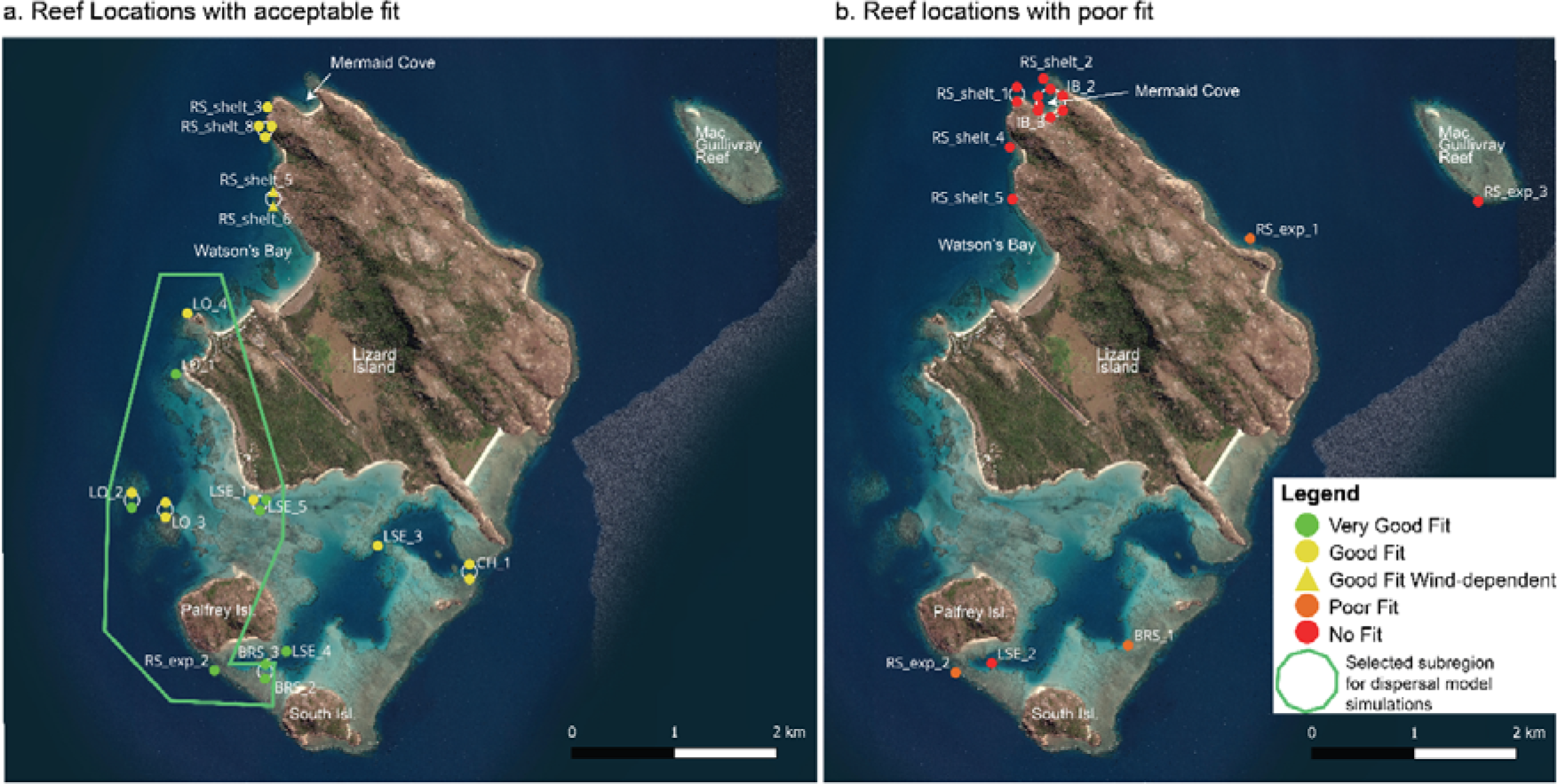
Map showing reef locations with acceptable fits (a) and poor fits (b) between modelled and observed current data based on findings from Table 1. The green polygon in (a) shows the subregion in the model domain particles dispersal ensuring best representation of real-world hydrodynamic conditions on the reef.

**Table 1:** Results from the validation analysis below 2 meters depth, comparing modelled current data with tiltmeter current data are several reefs during the study timeframe. Coloured cells highlight the strength of the linear relationship (R^2^) between modelled and observed velocity vectors (color gradient from green = good to red = poor). The fit category summarizes the overall representation of local current conditions by the model after detailed visual interpretation (Fig. S2-8) and expert knowledge explanations.

| Reef Location ID | Month Year | Depth | Reef Habitat | n | R<sup>2</sup><sub>u</sub> | R<sup>2</sup><sub>v</sub> | Fit Category | Expert knowledge explanations - more details in Figures S2-8 |
| --- | --- | --- | --- | --- | --- | --- | --- | --- |
| BRS_2 | 2021-11 | 1.4 | Back Reef Slope | 135 | 0.660 | 0.797 | Very Good Fit |  |
| BRS_3 | 2022-12 | 2.3 | Back Reef Slope | 289 | 0.615 | 0.051 | Very Good Fit | a small error in direction resulting in big error in v, but dominant direction is u for ebb and flow (Fig. S4) |
| LO_1 | 2021-08 | 5.1 | Lagoon Open | 70 | 0.361 | 0.632 | Very Good Fit |  |
| LO_2 | 2023-02 | 3.5 | Lagoon Open | 214 | 0.073 | 0.767 | Very Good Fit | a small error in direction resulting in big error in v, but dominant direction is u for ebb and flow (Fig. S5) |
| LSE_1 | 2021-08 | 3.7 | Lagoon SE | 70 | 0.859 | 0.363 | Very Good Fit |  |
| LSE_4 | 2021-11 | 2.6 | Lagoon SE | 137 | 0.661 | 0.402 | Very Good Fit |  |
| LSE_5 | 2021-11 | 1.7 | Lagoon SE | 137 | 0.548 | 0.014 | Very Good Fit | very good direction but with direction peaks centred at 90 and 270, meaning a very small error in direction resulting in big error in v, but in reality, the current fit is extremely good- the current is east-west ebb-flow tidal and dominated by u (Fig. S3) |
| RS_exp_2 | 2023-02 | 3.9 | Outer Reef Slope | 215 | 0.523 | 0.524 | Very Good Fit |  |
| CH_1 | 2021-08 | 8.0 | Channel | 70 | 0.782 | 0.038 | Good Fit | Lag effect (Fig. S5,S6), however inaccurate in predicting the correct time of weak current flow |
| CH_1 | 2022-12 | 3.4 | Channel | 239 | 0.611 | 0.178 | Good Fit |  |
| LO_2 | 2021-08 | 4.8 | Lagoon_Open | 70 | 0.590 | 0.419 | Good Fit |  |
| LO_3 | 2021-11 | 2.5 | Lagoon_Open | 137 | 0.460 | 0.249 | Good Fit |  |
| LO_3 | 2022-12 | 3.3 | Lagoon_Open | 289 | 0.434 | 0.196 | Good Fit |  |
| LO_4 | 2023-02 | 3.5 | Lagoon_Open | 210 | 0.394 | 0.303 | Good Fit |  |
| LSE_3 | 2021-08 | 7.9 | Lagoon_SE | 70 | 0.199 | 0.510 | Good Fit |  |
| LSE_5 | 2022-12 | 2.4 | Lagoon_SE | 239 | 0.417 | 0.288 | Good Fit |  |
| RS_shelt_8 | 2022-09 | 4.3 | Outer Reef Slope | 287 | 0.003 | 0.469 | Good Fit | Great representation of slack conditions (Fig. S7) |
| RS_shelt_8 | 2022-12 | 4.3 | Outer Reef Slope | 315 | 0.157 | 0.482 | Good Fit |  |
| RS_shelt_8 | 2023-02 | 4.3 | Outer Reef Slope | 210 | 0.249 | 0.579 | Good Fit |  |
| RS_shelt_3 | 2022-09 | 4.7 | Outer Reef Slope | 286 | 0.196 | 0.049 | Good Fit | But speed amplitude not always well represented (Fig. S7) |
| RS_shelt_5 | 2022-06 | 6.0 | Outer Reef Slope | 141 | 0.001 | 0.380 | Good Fit_wind | Better fit when Northward current, likely driven by southerly winds. Modelled current has a southward peak which is not apparent in reality and could be caused by a sheltering effect from northerly wind (Fig. S7) |
| RS_shelt_6 | 2022-06 | 2.9 | Outer Reef Slope | 141 | 0.045 | 0.231 | Good Fit_wind |  |
| RS_shelt_7 | 2022-06 | 7.3 | Outer Reef Slope | 141 | 0.000 | 0.326 | Good Fit_wind |  |
| BRS_1 | 2021-08 | 1.6 | Back Reef Slope | 70 | 0.004 | 0.714 | Poor Fit | Out of phase and amplitude issue, likely missing topography effect (Fig. S4,S8) |
| RS_exp_1 | 2021-08 | 4.7 | Outer Reef Slope | 70 | 0.106 | 0.042 | Poor Fit |  |
| RS_exp_2 | 2022-12 | 3.7 | Outer Reef Slope | 225 | 0.174 | 0.048 | Poor Fit | Out of phase and amplitude issue, likely missing topography effect (Fig. S4,S8), however, slack current conditions properties are represented |
| IB_1 | 2021-11 | 2.1 | Inner Bay | 136 | 0.133 | 0.229 | No Fit | Bay effect (Fig. S1) |
| IB_2 | 2021-11 | 2.6 | Inner Bay | 136 | 0.134 | 0.001 | No Fit |  |
| IB_2 | 2022-06 | 2.2 | Inner Bay | 139 | 0.035 | 0.061 | No Fit |  |
| IB_2 | 2022-09 | 2.3 | Inner Bay | 214 | 0.147 | 0.208 | No Fit |  |
| IB_2 | 2022-12 | 2.3 | Inner Bay | 287 | 0.028 | 0.006 | No Fit |  |
| IB_3 | 2022-06 | 5.4 | Inner Bay | 139 | 0.088 | 0.025 | No Fit |  |
| IB_4 | 2022-06 | 6.2 | Inner Bay | 139 | 0.018 | 0.031 | No Fit |  |
| LSE_2 | 2021-08 | 9.4 | Lagoon SE | 70 | 0.050 | 0.014 | No Fit | Lagoon local shelter effect (Fig. S3) |
| RS_exp_3 | 2022-12 | 4.5 | Outer Reef Slope | 223 | 0.011 | 0.018 | No Fit | Localized reef shelter effect (Fig. S8) |
| RS_shelt_1 | 2021-08 | 4.2 | Outer Reef Slope | 70 | 0.181 | 0.008 | No Fit | Model represents tides but not localized conditions (Fig. S7) |
| RS_shelt_1 | 2021-11 | 5.6 | Outer Reef Slope | 158 | 0.056 | 0.152 | No Fit |  |
| RS_shelt_2 | 2022-09 | 3.7 | Outer Reef Slope | 214 | 0.003 | 0.001 | No Fit |  |
| RS_shelt_4 | 2022-12 | 3.4 | Outer Reef Slope | 239 | 0.024 | 0.272 | No Fit | Localized land/reef shelter effect (Fig. S7) |
| RS_shelt_5 | 2021-11 | 2.9 | Outer Reef Slope | 136 | 0.007 | 0.424 | No Fit | Model represents tides but not localized conditions (Fig. S7) |

The model performed well in the eastern and south-eastern part of the domain, from Palfrey and South islands, to the central eastern ‘open lagoon’ habitat, to Osprey reef (Table 1, Fig. 4, Fig. S2-8). These areas encompass a variety of habitats: open lagoon, outer reef slope, back reef slope and channel. Reef locations that were dominated by the tidal and wind-driven circulation performed the best, for all habitats. Reef locations that were close to local reef and channels that deformed or delayed the tide propagation displayed less optimal predictions, sometimes with a lag effect (e.g. CH_1, LO_3). Reef locations that were sheltered by a local reef nearby performed poorly, with the model overpredicting the amplitude of the flow (e.g. BRS_1, LS_2, RS_exp_3) (Figs. S2-8).

The model performed poorly in the north of the domain, especially in the inner bay of Mermaid Cove (e.g. IB_1-4), with extremely weak current modelled inside the bay (Fig. S2). Outside the bay, model performance of predicted current speed improved at the points RS_shelt_1 and RS_Shelt 2 (Fig. S7) but the local deformation of the tide and the effect of headlands on circulation were not well represented. Along Watson’s Bay northern sheltered reef slope, modelled current was overall too weak but represented well the tidal phase and direction of the ebb and slack flow currents (RS_shelt_3, RS_shelt_8, Fig. S7). Modelled local current conditions further south at sites RS_shelt_5-7 (Fig. S7) was more accurate during northward flow but was unrealistic during southward flow. This behaviour is probably due to a sheltering effect of the island from the wind that is not accounted for in the atmospheric forcing.

As a result, dispersal simulations during the larval delivery phase, which occurs locally on shallow reefs deeper than 2 m were exclusively conducted at sites located within the subregion of the model domain that exhibited acceptable model validation findings (Fig. 4a).

### Predicting the residency of larvae over reefs during larval delivery phase

Particle Residency Time (PRT) was estimated at the 25 reef locations across the subregion where the model demonstrated acceptable validation. PRT exhibited considerable variation among sites and delivery timings (GLMM-POISSON, significant additive effect P < 0.001) (Fig. 5, S12). This finding indicates that (1) some sites such as in between the Palfrey and South Island (i.e., Sites 33-35) have overall better hydrodynamic properties at retaining particles than others and (2) overall, there were specific times of the day that lead to higher PRT (Figure S12). These findings suggest that when the optimal timing for high particle residency at a given location is identified, the potential for larvae to be retained in high concentrations from free larval releases could replicate conditions of a 3- to up to 12-hour larval settlement period in a mesh net enclosure (*sensu* dela Cruz and Harrison 2017, Harrison et al. 2021) but at much larger hectare scales (Fig. 5, S12). Among the 25 studied delivery sites, 16 sites exhibited at least one occurrence during the studied delivery timeframe when PRT was ≥5 hours within one hectare of the delivery site following delivery (Figure 5). Such occurrences were more frequent at sites located on the outer reef slope between Palfrey and South islands (i.e., Site 33-35). These reef locations specifically displayed a high frequency of long PRT make them ideal candidates for larval releases (Fig. 5).

**Figure 5:**
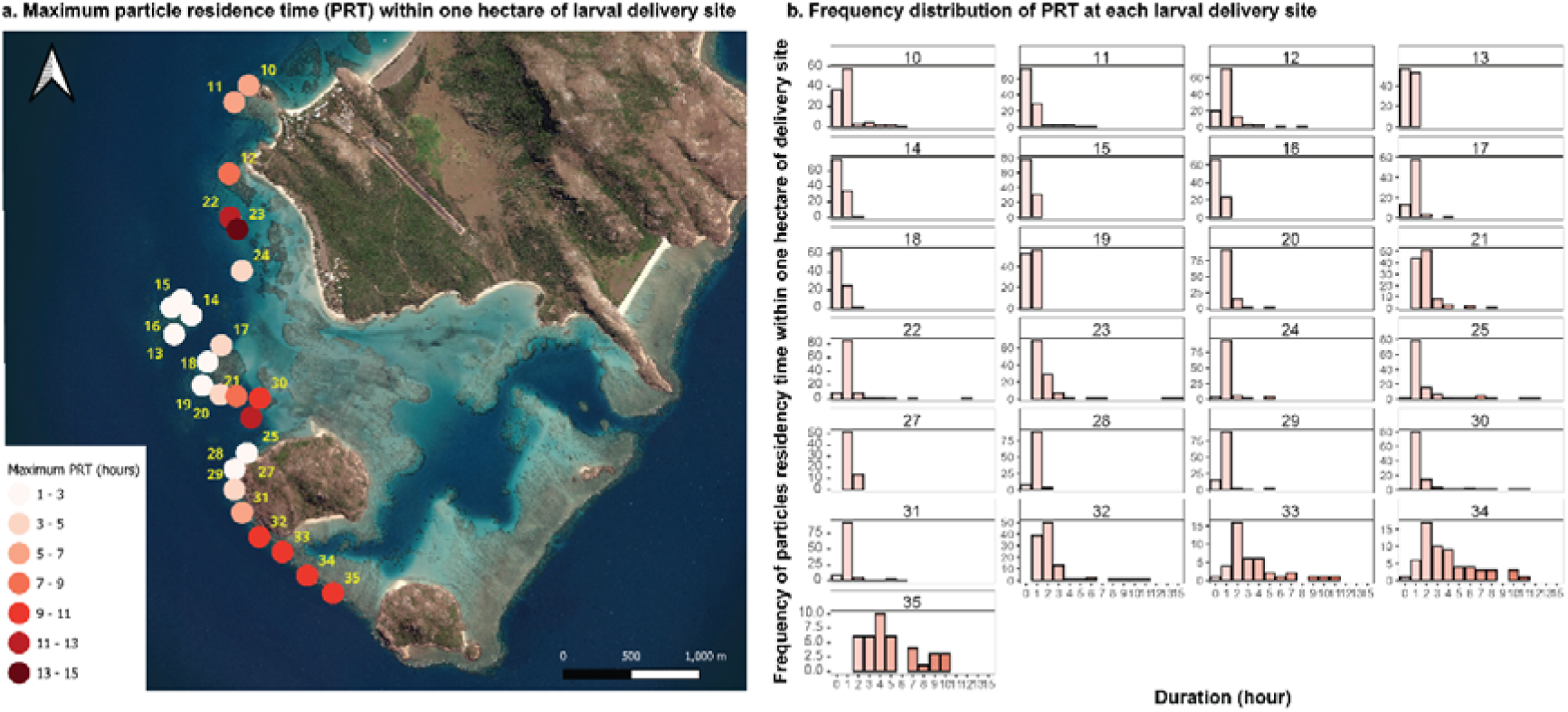
Bubble plot showing site with the maximum particles’ residency time within one hectare of the 20 delivery sites showing PRT _≥_5 hours (red colour scale) (a) and the frequency distribution of PRT at each of the delivery locations during the delivery timeframes in both 2021 and 2022. Number shows the locations ID. Sites with PRT lower than 5 were excluded.

The reefs between the two Palfrey Islands were occasionally observed to be sheltered from the dominant current conditions by topographic features such as Palfrey Island reefs. This occurred intermittently during time periods when the dominant current was flowing more southward (Figure 6.b.) as opposed to south-eastward for instance (Figure 6.a.). In addition, when the current oscillated from south eastward and north westward, small-scale eddies formed along the reef (Figure 6.c.). These types of conditions led to weaker current conditions on the reef south-east of Palfrey Island and promoted retention

**Figure 6:**
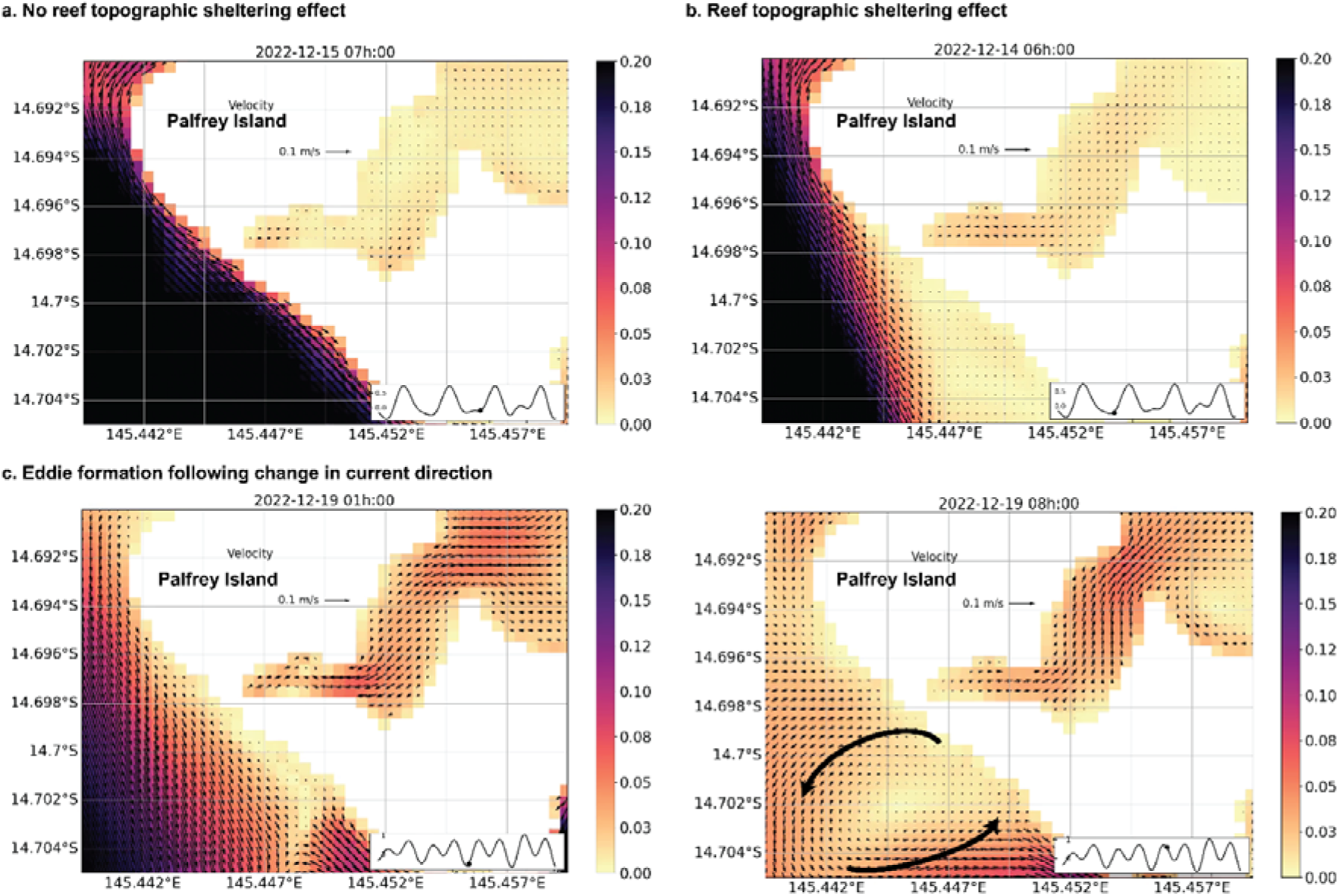
Snapshot of hydrodynamic conditions at 3.6 m modelled depth during delivery 2022 delivery time period, showing no reef topographic sheltering effect (a), topographic sheltering effect from Palfrey Island reef to the southeastward reefs (b) and eddie formation following a change in current direction (c). White cells represent reefs shallower than 3.6 m or land; black arrows show the current direction and coloured cells’ current speed in m.s-1.

### In situ *current speed conditions on the reef during larval delivery phase:* a posteriori Investigation

Current speed on the reef during larval delivery conditions (i.e., 5 to 10 days following a coral spawning event) at depths ranging from 2 to 5 m was explored *a posteriori* of reef intervention. Current speed was highly variable among the 17-time series (GLM, P<0.001) (Fig. 7.a., S2-S8). It was the lowest overall in the sheltered inner bay of Mermaid cove and two outer reef locations in the northern part of the island in December 2022 (Fig 7.a), with median current speeds lower than 0.01 m.s^-1^. Highest current regimes were observed in the lagoonal habitat with median current speeds >0.055 m.s^-1^ and maximum current speeds >0.2 m.s^-1^.

**Figure 7:**
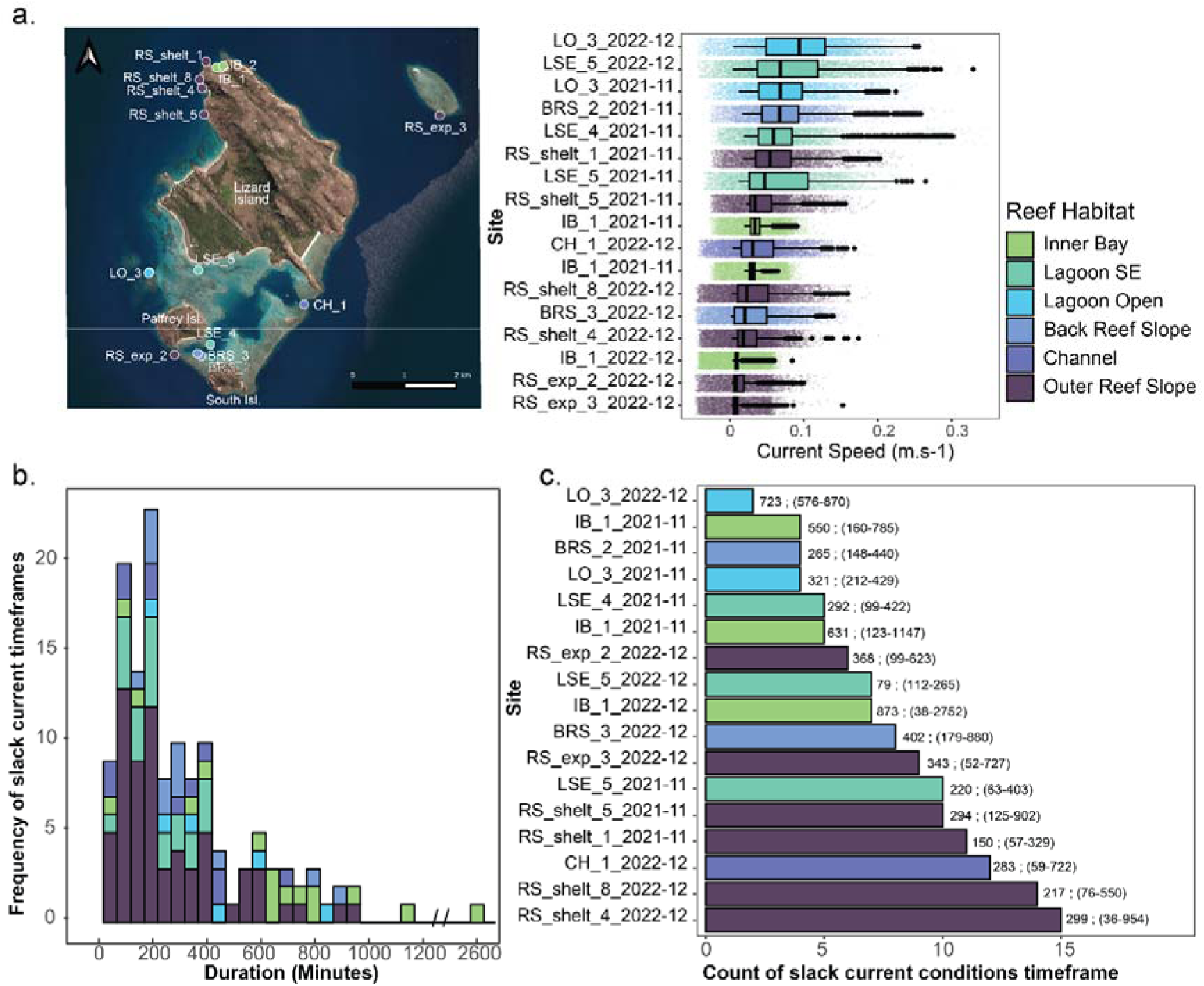
Overall current speed profiles at 17 sites encompassing 6 reef habitats in 2021 and 2022 (a), the frequency of occurrence of slack current conditions and their associated duration (b), and the count of slack current conditions at each of the study locations (c) number on the right to the bar shows the average and range of duration.

Slack current conditions (detected as timeframes of slow current speed generally < 0.022 m s-2) conformed to a log-normal long-tail distribution which ranged from 36 mins to 45 hours with a peak value of ∼ 3hours (Fig 7.b). The tail values are only occurring in the sheltered inner bay of Mermaid Cove. The most frequent slack current conditions lasted between 1.5 to 3.5 hours (Fig 7.b). The duration of slack current conditions was variable and significantly different among study sites (GLM NB P <0.001), but there were no clear patterns among habitats (Fig. 7.b). There were occurrences of extended slack current conditions lasting over 400 min (i.e., 6.5 hours) on reefs in all habitats during the delivery timeframe. Reefs that displayed frequent occurrences of long slack current conditions such as Mermaid Cove (IB_1, IB_2), along the reef slope of North Watson Bay (RS_shelt_8,4,5) or the outer reef slope between Palfrey and South Island (RS_exp_2), would make ideal candidate areas for deployment of coral larvae (Fig. 7.c). In contrast, reefs displaying few slack current conditions over the same timeframes such as reefs near Big Vickies (LO_3) are riskier to select for larval releases as delivery plumes will likely disperse very fast from the delivery source (Fig. 7.c). The timing of low and high tides was a poor predictor of slack current conditions and therefore should not be used to estimate slack current conditions (Fig. S13).

## Discussion

The main challenge in upscaling larval-based restoration of benthic marine invertebrates lies in identifying where to collect gametes and developing embryos *in situ* following reproductive events and when and where to release competent larvae to significantly increase local larval settlement and recruitment on target reef areas. This requires the integration of understandings of both fine-scale propagule dispersal and early developmental biology and behaviour of propagules. This study aimed to evaluate the application and accuracy of 3D reef-scale dispersal modelling in predicting the transport of coral spawn and larvae in the context of reef restoration interventions. Our findings show that near the surface, during propagules’ buoyant phase, the model accurately predicted the direction of coral spawn dispersal following spawning events and the occurrence of slick convergence zones allowing us to identify optimal locations for gametes and embryos collection. Below the surface and at the reef benthos scale (∼30 m-50 m), we found that the model occasionally failed to capture local-scale variability in current speed and direction (ie. at 42% of studied locations). This limitation was particularly evident at sites with significant sheltering effects not accounted for due to the bathymetric resolution of the model (30 m). However, at locations where the model predictions were accurate, we were able to predict with confidence the maximum residency times of propagule during the larval delivery phase (ie. neutrally buoyant phase) within one hectare of restoration sites. Our results highlight that long particle residence time were infrequent, but could last 5-15 hours at 15 of the 25 studied location locations when particles were released at specific days and times. *In situ* current observations demonstrated similar time range in slack current conditions (6.5 to 15 hours) during delivery timeframes in all reef habitats. Thus, our study has identified a workflow to identify the existence of these optimal, high retention periods that minimise dispersal away from, and maximize larval retention within, restoration sites without the use of equipment to constrain larvae as has been done in previous studies (Heyward et al. 2002, dela Cruz and Harrison 2017, 2020, Harrison et al. 2021). Overall, our findings highlight the potential of harnessing natural conditions to operationalise the mass collection and the mass settlement of released propagules for larval-based coral restoration efforts at larger hectare scales.

### Predicting spawn slick convergence zones for coral larval collection efforts

To maximize the collection of coral spawn following key coral spawning nights, modelled dispersal of particles near the surface was conducted at the scale of the island to identify convergence zone of high concentrations of particles. Most of the modelled particle tracks exhibited an accurate direction with a median error of 42 degrees when compared to drifters, however, modelled dispersal displayed overall slower drifts, leading to median errors in distance between particles and drifters of 323 meters. However, at the scale of the island, the model was able to predict convergence zone of spawn slicks that were witnessed during coral spawn collection at nights, such as north-westward and westward from Palfrey Island following key spawning events (authors pers obs, Fig. 3). These modelled convergence zones of highly concentrated coral spawn were more evident when the surface wind drag on particles was excluded from the simulations. Accurately identifying these convergence zones is important for larval based restoration.

Recent research on the distribution of floating materials in the ocean surface, such as oil or plastics (reviewed in Van Sebille et al., 2020), indicates that surface convergence zones at similar scales to this study are primarily driven by the combination of ocean currents, tidally generated eddies, windage, fronts, and Langmuir circulation (Wolanski and Hamner 1988, Wolanski et al. 1989). This phenomenon leads to the accumulation of debris/particles at locations where Langmuir cells converge (Dethleff et al. 2009). For positively buoyant particles such as coral spawn and developing embryos, Bees et al. (1998) described two studied behaviours. Some particles are trapped in closed orbits at a distance below the surface, known as the “Stommel retention zone”, while others accumulate at the convergence line at the surface. Analysis of forces acting on particles, including the buoyancy force and dynamic pressure force reveals that particles recirculate, follow different retention trajectories based on particle size and density (Dethleff et al. 2009, Chubarenko et al. 2016). Coral eggs and developing embryos are typically smaller (<0.5mm) than the smallest microplastics studied in this context (0.5 to 5mm) (Chubarenko et al. 2016), and are sticky and often clumped to each other to form extensive surface slicks (Oliver and Willis 1987) due to their hydrophobic quality caused by their high lipid content (Arai et al. 1993). *In situ* observations of coral spawn slicks have shown that their occurrence is closely associated with surface oceanographic features including wakes, eddies behind reefs, boundary mixing (where water of different density is mixed near sloping reefs), tidal jets (where currents are squeezed between two reefs), and lastly windage (reviewed in Pattiaratchi 1994, Wolanski et al. 2024).

With a 30-50m resolution, the hydrodynamical model can represent dynamical processes resulting in convergence zones at a scale of 100s of meters. However, local windage effect relies on accurate wind forcing datasets. With ACCESS-G resolution of 12 km, the windage effect is likely misrepresented in the current dispersal experiments. Moreover, the current dispersal experiments are performed in 2D. Wind forcing is known to influence the vertical mixing rate of coral spawn (Wolanski et al. 1989). *In situ* observations show that during windy conditions developing embryos can be mixed throughout the top 10 m; but in low wind conditions embryos aggregate at the surface (Oliver and Willis 1987, Wolanski et al. 1989, Willis and Oliver 1990, Pattiaratchi 1994) (Figure 3.b). While the vertical mixing effect due to the wind is represented in the hydrodynamical model, this effect is lost when using a 2D current field for the dispersal modelling. Therefore, improving our understanding on the physical dispersal properties of coral spawn, which differ from microplastics or oil slicks (Wolanski et al. 1989, Willis and Oliver 1990), and how local wind conditions influence it, is critical for accurately modelling its buoyant phase (Pattiaratchi 1994).

### Predicting the local variability in current conditions at reef-scale for larval deployment

Modelling hydrodynamic conditions within reef scales (1-100’s of metres) is extremely complex (Swearer et al. 2019, Wolanski et al. 2024) but highly applicable to propagule collection and deployment during restoration interventions. Forecasting particle retention before deploying larvae onto reefs aids in determining the optimal location, day, and time for delivery, thereby maximizing retention over damaged reefs, promoting mass settlement, and enhancing intervention success. Our validation results show a varied performance of the model in forecasting accurate local hydrodynamic conditions on reefs. Firstly, when tides and wind-driven currents dominated the current signal, the model exhibited its best performance, applicable across all studied habitats, resulting in more than half of the validation time series displaying good to very good validation results. Secondly, in small, protected bays like Mermaid Cove (sites IB_1,2, Fig. S2), the model resolution was inadequate, resulting in slow or negligible modelled currents. *In situ* current observations in this habitat showed weak currents (< 0.02 m/s for long periods), but with magnitudes higher than modelled currents, resulting in a small but apparent deviation between observed and modelled currents. Thirdly, the model struggled to accurately represent conditions where local reefs or structures created a shelter effect from dominant current flow. For instance, persistent strong tidal currents were simulated by the model despite observations indicating weak non-tidal currents at site RS_shelt_5 (see Fig. S7). Despite the ∼30 m resolution, the inability of the model to fully capture reef-induced circulation or sheltering effects at a small scale is evident in complex topographic settings, as shown in the large differences in flow speeds and direction between topographic highs and lows from the instrument data quantified within the reef area of a model grid cell (Fig. S11). Lastly, certain studied reef locations were partially influenced by local reef structures, resulting in instances where the tidal signal was distorted or delayed (see Fig. S6). While the performance of the model at these locations was acceptable under specific conditions, improvements could be made by refining the bathymetry, particularly in areas with significant observed sheltering effects.

At locations where the model predictions were acceptable, we modelled the residence time of particles during 24 hours following their delivery onto 25 reefs during the entire ‘day time’ larval delivery timeframes in both 2021 and 2022. Our simulations show that high particle residency periods, ranging from 5 to 15 h, occurred on at least one occasion at 60% of studied sites (ie. 15 of 25). Identifying the location of such reefs and the start time of prime retention windows ahead of larval deployment day(s) would help to inform decisions on when to release competent larvae to maximize their retention over the damaged reef. Among the 25 locations studied, the most frequent instances of long particle residency time (>5 h) within a hectare of delivery location were observed along the reef slope between Palfrey and South Island (Fig. 5). Although similar PRT occurred elsewhere, they were less frequent (Fig. 5.b.). Through an examination of modelled hydrodynamic conditions and particles trajectories, primary retention mechanisms that led to particle aggregation and high residency near the delivery location were identified. These local retention mechanisms are purely physical, directly related to transport by local hydrodynamic conditions, which are known to occur on reefs (Wolanski et al. 1996, Sponaugle et al. 2002, Cetina-Heredia and Connolly 2011). Firstly, tidal currents plays a crucial role in particle retention by creating back-and-forth displacement and slack current conditions on reefs (Wolanski et al. 2024). Tidal currents, cycling on a 6-hour interval in this study, can drive the retention of particles over the restoration site during slack conditions or, outside of slack conditions, transport particles over the same location up to three times over a 24-hour timeframe (e.g., Fig. S14a). Secondly, one of the other observed retention mechanisms observed along the outer reef slope between Palfrey and South Island, was a topographic sheltering effect (i.e., island wake effect (Wolanski et al. 1984, Pattiaratchi et al. 1987)), where the dominant current detached from the coastline in the lee from Palfrey Island and reef, created weak current conditions for several hours along most of the reef South of Palfrey Island (Figure 7a). Thirdly, when the dominant current changed to the opposite direction (primarily driven by tides here), the formation of a small-scale Eddie measuring ∼500 m across was modelled. The formation of this small eddy, generating oscillating current transported the larval plume southwestward, northeastward, and southeastward, resulted in small aggregation patterns of particles near the delivery site (Fig. S14.b). This phenomenon was shown in other studies to trap larvae locally from minutes to hours (Wolanski et al. 1996, Delandmeter et al. 2017) and is discussed in the recent study by Philipps and Bellwood (2024) as a potential driver for fast natural coral recruitment rate along that reef slope. Lastly, the presence of shallow reefs (< 3 m depth) between the two islands acted as a barrier to dispersal > 3 m depth in a northeast-southwest direction, blocking dispersal in and out of the lagoon at that depth (further discussed below in the *Study limitations* section).

### In situ *slack current conditions on reefs during larval deployment: making sense of modelled particle residency*

By exploring *in situ* current datasets *a posteriori* of larval deployment, we found that slack current conditions on the reef ranged from 36 mins to 45 hours; the latter only occurring in the sheltered inner bay of Mermaid Cove (Fig. 7). These *in situ* observations shared some similarity with modelled PRT. For instance, during several delivery scenarios in December 2022 (Fig. 5), modelled PRT along the reef slope between Palfrey and South Islands was lasting up to 12 hours, matching the long slack current conditions observed at a nearby location (RS_exp_2, Fig. S8). Throughout December 2022, *in situ* current speeds on the reef were generally slow, with extended periods of slack currents lasting up to 12 hours at a time (RS_exp_2, Fig. S8).

Several additional reef areas exhibited frequent occurrences of similar long slack current conditions and were mainly located in sheltered inner bay and on reef slopes (Fig. 7). These locations included the sheltered inner bay of Mermaid Cove (IB_1, IB_2), which displayed very long periods of slack and slow current, typical of this type of reef habitat (e.g. Ouillon et al. 2010, Golbuu et al. 2016). The reef location ‘RS_exp_3’, situated southwest of MacGillivray reef, experienced several consecutive days of very weak currents (Figs. 7, S8, S13) likely caused by an island wake process (Wolanski et al. 1984). Similarly, along the reef slope of North Watson Bay (RS_shelt_8, 4, 5), slack current conditions persisted for an average of 4.5 hours but could last for a maximum duration of 15 hours (Fig. 7). Combinedly, these observed conditions at these locations are ideal for the release of coral larvae and suggest that modelled PRT findings reaching 12-15 hr may likely be occurring at some reefs, as also demonstrated in Shedrawi et al. (2017), providing opportunities to deliver larvae during optimal conditions, maximizing local larval retention for reef restoration efforts.

### Study limitations and improving fine-scale dispersal modelling for larval-based restoration

The dispersal model in this study offers valuable insights into larval transport dynamics, yet several limitations were acknowledged from the beginning of the study. These limitations merit discussion for future improvement in modelling hydrodynamics at scales less than 30-50 m around complex shallow benthic habitats such as coral reefs. Identified model limits include local-scale wind conditions, island shadow effect (Critchell and Lambrechts 2016), uniform bottom friction inadequately representing diverse substrata types across coral reef habitats (Sous et al. 2022), and coarse bathymetry maps (although the finest possible for this type of study). All of these could influence the accurate representation of circulation phenomena such as small eddies, island wakes, and topographic shelter effects occurring at fine scales. For example, discrepancies arose in predicting local current conditions at sites influenced by sheltering effects or near complex reef structures. Addressing this limitation necessitates finer bathymetric data that can be collected through multibeam sonar (e.g. Zhi et al. 2014, Colin et al. 2019). The detailed imagery obtained from shallow multibeam sonar would capture substrate roughness and benthic habitat complexity, likely facilitating the resolution of reef-induced circulation and boundary layer dynamics currently absent in hydrodynamic model predictions. While this improvement may be advantageous, it would also trade-off with increased processing times, and therefore requiring careful consideration depending on objectives.

Using the particle tracking model component CONNIE, neutrally buoyant particles remained bound to their dispersal depth in the 24-hr dispersal scenarios, which was 3.6 m throughout the 24-hour dispersal in the larval deployment simulations of this study. This was a key limitation of the particle tracking model component. Modelling particles that are fully dispersed in 3D following their release depth is an important consideration for future fine-scale dispersal modelling (e.g. Takeyasu et al. 2023). This improvement would prevent particles from getting stuck between depth layers and would allow investigation into whether, for example, particles remain at the same depth where weak currents are found or if upwelling or downwelling currents could transport them to shallower or deeper reef areas.

Integrating additional observational data, including high-resolution current measurements and remotely sensed data, is crucial for advancing model development and strengthening confidence in predictions as demonstrated in this study. Increasing the spatial coverage and duration of *in situ* current data collection would strengthen future studies. Deploying expensive instruments for long periods of time is high-risk, yet longer time series of *in situ* current data are essential for capturing the full spectrum of hydrodynamic variability at local scales and improving model performance. In addition, improving the tracking of surface spawn slicks is also suggested. Drifters used in this study capture current flow 1 m below the surface and may not adequately measure the drift dynamics of slicks at the surface which are highly subjected to wind drag effects (Morey et al. 2018). Exploring innovative techniques, such as utilizing specialized drifters designed to track the top surface currents more effectively, such as in Sutherland et al. (2020), or employing night-time drones for high-resolution observations of coral spawn slicks (Mohsan et al. 2023) could offer valuable insights into spawn slick transport at fine temporal scales. These advances in observational methods hold promise for enhancing our understanding of larval dispersal and improving the predictive capability of dispersal models at reef scales, not only for coral reef restoration efforts but also for broader scale dispersal models that lead to improved connectivity estimates.

### Conclusion

This study described and tested the use of a fine-scale dispersal modelling to inform effective strategies for larval-based reef restoration. By integrating *in situ* observations, field data, and dispersal simulations, we highlight that the scalability of coral spawn collection and larval delivery is possible. This was characterised by predictions of convergence zones of coral spawn slicks often observed following major spawning events, and optimal timeframes of long particle residency over damaged reefs to encourage local settlement ‘en masse’. Our findings underscore the need for further improvements in modelling accuracy to better predict hydrodynamic conditions at the reef scale, predominantly relating to finer-scale bathymetry mapping. Ultimately, advancing our understanding of larval dispersal processes is essential for successful reef restoration efforts and the preservation of these vulnerable ecosystems in the face of global climate pressures.

## Acknowledgements

The authors acknowledge the Traditional Owners of Lizard Island on which this research was conducted, the Dingaal, Thaanil-Warra and Ngurruumungu Aboriginal people, and their ongoing connection to the land and sea. We sincerely thank John Andrewartha for creating the original hydrodynamic model and early advice into the study. We thank Mark Tonks and Robert Mason for their help collecting hydrodynamic data in August-September 2021 and Bec Gorton for her expertise on using CONNIE. We also thank staff of the Lizard Island Research Station for providing support during field work activities. This research was funded by the Moving Corals subprogram of the Reef Restoration and Adaptation Program (RRAP) to PH and CD. The Reef Restoration and Adaptation is funded by the partnership between the Australian Government’s Reef Trust and the Great Barrier Reef Foundation. The funders had no role in study design, data collection and analysis, decision to publish, or preparation of the manuscript. This study was conducted under GBRMPA permit no. G20/44511.1.

## Author Contributions

Marine Gouezo, Clothilde Langlais and Christopher Doropoulos conceived the study idea and designed methodology; Clothilde Langlais and Jack Beardsley created the model; Marine Gouezo, Clothilde Langlais and Jack Beardsley ran the dispersal simulations; Marine Gouezo, Damian Thomson, George Roff, Peter Harrison and Christopher Doropoulos collected *in situ* hydrodynamic data and other observations; George Roff modelled the relative relationship between coral larvae density and live coral cover; Marine Gouezo and Clothilde Langlais extracted data from dispersal modelling simulations, Marine Gouezo ran all statistical analyses and led the draft of the manuscript. All authors contributed critically to the manuscript drafts and gave final approval for publication

## References

AIMS. 2023. AIMS Long-term Monitoring Program: Crown-of-thorns starfish and benthos Manta Tow Data (Great Barrier Reef).

Allen Coral Atlas. 2022. Imagery, maps and monitoring of the world’s tropical coral reefs.

Arai, I., M. Kato, A. Heyward, Y. Ikeda, T. Iizuka, and T. Maruyama. 1993. Lipid composition of positively buoyant eggs of reef building corals. Coral Reefs 12:71–75.

Asner, G. P., N. R. Vaughn, R. E. Martin, S. A. Foo, J. Heckler, B. J. Neilson, and J. M. Gove. 2022. Mapped coral mortality and refugia in an archipelago-scale marine heat wave. Proceedings of the National Academy of Sciences 119:e2123331119.

Australian Bureau of Meteorology (ABOM). 2017. Operational upgrades to OceanMAPS (BLUElink> ocean forecast system) – global ocean forecasting. Operations Bulletin 89.

Baird, M. E., K. A. Wild-Allen, J. Parslow, M. Mongin, B. Robson, J. Skerratt, F. Rizwi, M. Soja-Woźniak, E. Jones, and M. Herzfeld. 2020. CSIRO Environmental Modelling Suite (EMS): scientific description of the optical and biogeochemical models (vB3p0). Geoscientific Model Development 13:4503–4553.

Banaszak, A. T., K. L. Marhaver, M. W. Miller, A. C. Hartmann, R. Albright, M. Hagedorn, P. L. Harrison, K. R. W. Latijnhouwers, S. Mendoza Quiroz, V. Pizarro, and V. F. Chamberland. 2023. Applying coral breeding to reef restoration: best practices, knowledge gaps, and priority actions in a rapidly evolving field. Restoration Ecology 31:e13913.

Bayraktarov, E., M. I. Saunders, S. Abdullah, M. Mills, J. Beher, H. P. Possingham, P. J. Mumby, and C. E. Lovelock. 2016. The cost and feasibility of marine coastal restoration. Ecological Applications 26:1055–1074.

Beck, M. W., R. D. Brumbaugh, L. Airoldi, A. Carranza, L. D. Coen, C. Crawford, O. Defeo, G. J. Edgar, B. Hancock, and M. C. Kay. 2011. Oyster reefs at risk and recommendations for conservation, restoration, and management. Bioscience 61:107–116.

Bivand, R., T. Keitt, B. Rowlingson, E. Pebesma, M. Sumner, R. Hijmans, E. Rouault, and M. R. Bivand. 2015. Package ‘rgdal.’ Bindings for the Geospatial Data Abstraction Library. Available online: https://cran.r-project.org/web/packages/rgdal/index.html (accessed on 15 October 2017) 172.

Bode, M., J. M. Leis, L. B. Mason, D. H. Williamson, H. B. Harrison, S. Choukroun, and G. P. Jones. 2019. Successful validation of a larval dispersal model using genetic parentage data. PLOS Biology 17:e3000380.

Bode, M., O. Stewart, and S. M. Choukroun. 2024. Incorporating biophysical larval dispersal simulations into coral reef conservation decision-making. Pages 272–281 Oceanographic Processes of Coral Reefs. CRC Press.

Bolin, B., and H. Rodhe. 1972. A note on the concepts of age distribution and transit time in natural reservoirs. Tellus A 25.

Boschetti, F., R. C. Babcock, C. Doropoulos, D. P. Thomson, M. Feng, D. Slawinski, O. Berry, and M. A. Vanderklift. 2020. Setting priorities for conservation at the interface between ocean circulation, connectivity, and population dynamics. Ecological Applications 30.

Broadhurst, L. M., A. Lowe, D. J. Coates, S. A. Cunningham, M. McDonald, P. A. Vesk, and C. Yates. 2008. Seed supply for broadscale restoration: maximizing evolutionary potential. Evolutionary Applications 1:587–597.

Bruyère, O., M. Chauveau, R. Le Gendre, V. Liao, and S. Andréfouët. 2023. Larval dispersal of pearl oysters Pinctada margaritifera in the Gambier Islands (French Polynesia) and exploring options for adult restocking using in situ data and numerical modelling. Marine Pollution Bulletin 192:115059.

Cetina-Heredia, P., and S. R. Connolly. 2011. A simple approximation for larval retention around reefs. Coral Reefs 30:593–605.

Chubarenko, I., A. Bagaev, M. Zobkov, and E. Esiukova. 2016. On some physical and dynamical properties of microplastic particles in marine environment. Marine pollution bulletin 108:105–112.

Colin, P. L., T. S. Johnston, J. A. MacKinnon, C. Y. Ou, D. L. Rudnick, E. J. Terrill, S. J. Lindfield, and H. Batchelor. 2019. NGARAARD PINNACLE, PALAU. Oceanography 32:164–173.

Condie, S., M. Hepburn, and J. Mansbridge. 2012. Modelling and visualisation of connectivity on the Great Barrier Reef. Pages 9–13 Proceedings of the 12th International Coral Reef Symposium.

Connolly, S. R., and A. H. Baird. 2010. Estimating dispersal potential for marine larvae: dynamic models applied to scleractinian corals. Ecology 91:3572–3583.

Couto, N., J. Kohut, O. Schofield, M. Dinniman, and J. Graham. 2017. Pathways and retention times in a biologically productive canyon system on the West Antarctic Peninsula. Pages 1–8 OCEANS 2017-Anchorage. IEEE.

Critchell, K., and J. Lambrechts. 2016. Modelling accumulation of marine plastics in the coastal zone; what are the dominant physical processes? Estuarine, Coastal and Shelf Science 171:111–122.

dela Cruz, D. W., and P. L. Harrison. 2017. Enhanced larval supply and recruitment can replenish reef corals on degraded reefs. Scientific Reports 7:13985.

dela Cruz, D. W., and P. L. Harrison. 2020. Enhancing coral recruitment through assisted mass settlement of cultured coral larvae. PLOS ONE 15:e0242847.

Delandmeter, P., J. Lambrechts, G. O. Marmorino, V. Legat, E. Wolanski, J.-F. Remacle, W. Chen, and E. Deleersnijder. 2017. Submesoscale tidal eddies in the wake of coral islands and reefs: satellite data and numerical modelling. Ocean Dynamics 67:897– 913.

Dethleff, D., E. W. Kempema, R. Koch, and I. Chubarenko. 2009. On the helical flow of Langmuir circulation—approaching the process of suspension freezing. Cold Regions Science and Technology 56:50–57.

Doropoulos, C., and R. C. Babcock. 2018. Harnessing connectivity to facilitate coral restoration. Frontiers in Ecology and the Environment 16:558–559.

Doropoulos, C., J. Elzinga, R. ter Hofstede, M. van Koningsveld, and R. C. Babcock. 2019a. Optimizing industrial-scale coral reef restoration: comparing harvesting wild coral spawn slicks and transplanting gravid adult colonies: Industrial-scale coral reef restoration. Restoration Ecology.

Doropoulos, C., L. A. Gómez Lemos, K. Salee, M. J. McLaughlin, J. Tebben, M. Van Koningsveld, M. Feng, and R. C. Babcock. 2022. Limitations to coral recovery along an environmental stress gradient. Ecological Applications 32:e2558.

Doropoulos, C., and G. Roff. 2022. Colouring coral larvae for tracking dispersal. preprint, Ecology.

Doropoulos, C., F. Vons, J. Elzinga, R. ter Hofstede, K. Salee, M. van Koningsveld, and R. C. Babcock. 2019b. Testing industrial-scale coral restoration techniques: harvesting and culturing wild coral-spawn slicks. Frontiers in Marine Science 6:658.

Duarte, C. M., S. Agusti, E. Barbier, G. L. Britten, J. C. Castilla, J.-P. Gattuso, R. W. Fulweiler, T. P. Hughes, N. Knowlton, and C. E. Lovelock. 2020. Rebuilding marine life. Nature 580:39–51.

Dumas, F., R. Le Gendre, Y. Thomas, and S. Andréfouët. 2012. Tidal flushing and wind driven circulation of Ahe atoll lagoon (Tuamotu Archipelago, French Polynesia) from in situ observations and numerical modelling. Marine Pollution Bulletin 65:425–440.

Edwards, A., J. Guest, A. Heyward, R. Villanueva, M. Baria, I. Bollozos, and Y. Golbuu. 2015. Direct seeding of mass-cultured coral larvae is not an effective option for reef rehabilitation. Marine Ecology Progress Series 525:105–116.

Fox-Kemper, B., A. Adcroft, C. W. Böning, E. P. Chassignet, E. Curchitser, G. Danabasoglu, C. Eden, M. H. England, R. Gerdes, and R. J. Greatbatch. 2019. Challenges and prospects in ocean circulation models. Frontiers in Marine Science 6:65.

Fulton, C. J., and D. R. Bellwood. 2005. Wave-induced water motion and the functional implications for coral reef fish assemblages. Limnology and Oceanography 50:255– 264.

Golbuu, Y., M. Gouezo, H. Kurihara, L. Rehm, and E. Wolanski. 2016. Long-term isolation and local adaptation in Palau’s Nikko Bay help corals thrive in acidic waters. Coral Reefs.

Gouezo, M., E. Wolanski, K. Critchell, K. Fabricius, P. Harrison, Y. Golbuu, and C. Doropoulos. 2021. Modelled larval supply predicts coral population recovery potential following disturbance. Mar. Ecol. Prog. Ser. 661:127–145.

Griffies, S. M., A. J. Adcroft, H. Banks, C. W. Böning, E. P. Chassignet, G. Danabasoglu, S. Danilov, E. Deleersnijder, H. Drange, and M. England. 2009. Problems and prospects in large-scale ocean circulation models. Proceedings of OceanObs 9:410–431.

Halpern, B. S., K. A. Selkoe, F. Micheli, and C. V. Kappel. 2007. Evaluating and ranking the vulnerability of global marine ecosystems to anthropogenic threats. Conservation Biology 21:1301–1315.

Halpern, B. S., S. Walbridge, K. A. Selkoe, C. V. Kappel, F. Micheli, C. D’Agrosa, J. F. Bruno, K. S. Casey, C. Ebert, H. E. Fox, R. Fujita, D. Heinemann, H. S. Lenihan, E. M. P. Madin, M. T. Perry, E. R. Selig, M. Spalding, R. Steneck, and R. Watson. 2008. A Global Map of Human Impact on Marine Ecosystems. Science 319:948–952.

Harrison, P. L., R. C. Babcock, G. D. Bull, J. K. Oliver, C. C. Wallace, and B. L. Willis. 1984. Mass spawning in tropical reef corals. Science 223:1186–1189.

Harrison, P. L., K. A. Cameron, and P. C. Cabaitan. 2021. Increased Coral Larval Supply Enhances Recruitment for Coral and Fish Habitat Restoration. Frontiers in Marine Science:1786.

Harrison, P. L., and D. dela Cruz. 2022. Methods for restoring damaged reefs using coral larval restoration.

Hartig, F. 2017. DHARMa: residual diagnostics for hierarchical (multi-level/mixed) regression models. R package version 0.1 5.

Heyward, A. J., L. D. Smith, M. Rees, and S. N. Field. 2002. Enhancement of coral recruitment by in situ mass culture of coral larvae. Marine Ecology Progress Series 230:113–118.

Hock, K., N. H. Wolff, J. C. Ortiz, S. A. Condie, K. R. N. Anthony, P. G. Blackwell, and P. J. Mumby. 2017. Connectivity and systemic resilience of the Great Barrier Reef. PLOS Biology 15:e2003355.

Hoegh-Guldberg, O. 1999. Climate change, coral bleaching and the future of the world’s coral reefs. Marine and freshwater research 50:839–866.

Hudson, K., M. J. Oliver, J. Kohut, M. S. Dinniman, J. M. Klinck, C. Moffat, H. Statscewich, K. S. Bernard, and W. Fraser. 2021. A Recirculating Eddy Promotes Subsurface Particle Retention in an Antarctic Biological Hotspot. Journal of Geophysical Research: Oceans 126:e2021JC017304.

Hughes, T. P., J. T. Kerry, M. Álvarez-Noriega, J. G. Álvarez-Romero, K. D. Anderson, A. H. Baird, R. C. Babcock, M. Beger, D. R. Bellwood, R. Berkelmans, T. C. Bridge, I. R. Butler, M. Byrne, N. E. Cantin, S. Comeau, S. R. Connolly, G. S. Cumming, S. J. Dalton, G. Diaz-Pulido, C. M. Eakin, W. F. Figueira, J. P. Gilmour, H. B. Harrison, S. F. Heron, A. S. Hoey, J.-P. A. Hobbs, M. O. Hoogenboom, E. V. Kennedy, C. Kuo, J. M. Lough, R. J. Lowe, G. Liu, M. T. McCulloch, H. A. Malcolm, M. J. McWilliam, J. M. Pandolfi, R. J. Pears, M. S. Pratchett, V. Schoepf, T. Simpson, W. J. Skirving, B. Sommer, G. Torda, D. R. Wachenfeld, B. L. Willis, and S. K. Wilson. 2017. Global warming and recurrent mass bleaching of corals. Nature 543:373–377.

Johansen, J. L. 2014. Quantifying Water Flow within Aquatic Ecosystems Using Load Cell Sensors: A Profile of Currents Experienced by Coral Reef Organisms around Lizard Island, Great Barrier Reef, Australia. PLoS ONE 9:e83240.

Killick, R., and I. Eckley. 2014. changepoint: An R package for changepoint analysis. Journal of statistical software 58:1–19.

Largier, J. L. 2003. CONSIDERATIONS IN ESTIMATING LARVAL DISPERSAL DISTANCES FROM OCEANOGRAPHIC DATA. Ecological Applications 13:71– 89.

Madin, J. S., K. P. Black, and S. R. Connolly. 2006. Scaling water motion on coral reefs: from regional to organismal scales. Coral Reefs 25:635–644.

Magnusson, A., H. Skaug, A. Nielsen, C. Berg, K. Kristensen, M. Maechler, K. van Bentham, B. Bolker, M. Brooks, and M. M. Brooks. 2017. Package ‘glmmtmb.’ R Package Version 0.2. 0 25.

Mohsan, S. A. H., M. A. Khan, and Y. Y. Ghadi. 2023. Editorial on the Advances, Innovations and Applications of UAV Technology for Remote Sensing. MDPI.

Morey, S. L., N. Wienders, D. S. Dukhovskoy, and M. A. Bourassa. 2018. Measurement characteristics of near-surface currents from ultra-thin drifters, drogued drifters, and HF radar. Remote Sensing 10:1633.

Morgan, S. G., S. H. Miller, M. J. Robart, and J. L. Largier. 2018. Nearshore larval retention and cross-shelf migration of benthic crustaceans at an upwelling center. Frontiers in Marine Science 5:161.

Nickols, K. J., B. Gaylord, and J. L. Largier. 2012. The coastal boundary layer: predictable current structure decreases alongshore transport and alters scales of dispersal. Marine Ecology Progress Series 464:17–35.

Oliver, J. K., and B. L. Willis. 1987. Coral-spawn slicks in the Great Barrier Reef: preliminary observations. Marine Biology 94:521–529.

Ooms, J. 2014, March 12. The jsonlite Package: A Practical and Consistent Mapping Between JSON Data and R Objects. arXiv.

Orth, R. J., T. J. Carruthers, W. C. Dennison, C. M. Duarte, J. W. Fourqurean, K. L. Heck, A. R. Hughes, G. A. Kendrick, W. J. Kenworthy, and S. Olyarnik. 2006. A global crisis for seagrass ecosystems. Bioscience 56:987–996.

Orth, R., K. Moore, S. Marion, D. Wilcox, and D. Parrish. 2012. Seed addition facilitates eelgrass recovery in a coastal bay system. Marine Ecology Progress Series 448:177– 195.

Ouillon, S., P. Douillet, J.-P. Lefebvre, R. Le Gendre, A. Jouon, P. Bonneton, J.-M. Fernandez, C. Chevillon, O. Magand, and J. Lefèvre. 2010. Circulation and suspended sediment transport in a coral reef lagoon: The south-west lagoon of New Caledonia. Marine pollution bulletin 61:269–296.

Pattiaratchi, C. 1994. Physical Oceanograhic Aspects Of The Dispersal of Coral Spawn Slicks: A Review. Pages 89–105 in P. W. Sammarco and M. L. Heron, editors. Coastal and Estuarine Studies. American Geophysical Union, Washington, D. C.

Pattiaratchi, C., A. James, and M. Collins. 1987. Island wakes and headland eddies: A comparison between remotely sensed data and laboratory experiments. Journal of Geophysical Research: Oceans 92:783–794.

Pebesma, E. 2012. spacetime: Spatio-temporal data in R. Journal of statistical software 51:1– 30.

Pebesma, E., R. Bivand, M. E. Pebesma, S. RColorBrewer, and A. A. A. Collate. 2012. Package ‘sp.’ The Comprehensive R Archive Network.

Pebesma, E. J. 2018. Simple features for R: standardized support for spatial vector data. R J. 10:439.

Philipps, C. J., and D. R. Bellwood. 2024. The hydrodynamics of Lizard Island lagoon, Great Barrier Reef. Coral Reefs.

Puri, K., G. Dietachmayer, P. Steinle, M. Dix, L. Rikus, L. Logan, M. Naughton, C. Tingwell, Y. Xiao, and V. Barras. 2013. Implementation of the initial ACCESS numerical weather prediction system. Australian Meteorological and Oceanographic Journal 63:265–284.

QGIS Development Team. 2023. QGIS Geographic Information System. Open Source Geospatial Foundation Project.

R Development Core Team. 2023. R: A language and environment for statistical computing. Vienna, Austria.

Randall, C. J., A. P. Negri, K. M. Quigley, T. Foster, G. F. Ricardo, N. S. Webster, L. K. Bay, P. L. Harrison, R. C. Babcock, and A. J. Heyward. 2020. Sexual production of corals for reef restoration in the Anthropocene. Marine Ecology Progress Series 635:203–232.

Riginos, C., K. Hock, A. M. Matias, P. J. Mumby, M. J. van Oppen, and V. Lukoschek. 2019. Asymmetric dispersal is a critical element of concordance between biophysical dispersal models and spatial genetic structure in Great Barrier Reef corals. Diversity and Distributions 25:1684–1696.

Rinkevich, B. 1995. Restoration strategies for coral reefs damaged by recreational activities: the use of sexual and asexual recruits. Restoration Ecology 3:241–251.

Roelfsema, C. M., M. Saunders, R. Canto, J. X. Leon, S. R. Phinn, and S. Hamylton. 2014. Habitat Map for Lizard Island reef, Australia derived from a photo-transect survey field data collected in December 2011 and September/October 2012.

Roff, G. 2023. Spatially explicit modelling of coral spawning on the Great Barrier Reef.

Santos, L. A., K. R. Ferreira, G. R. de Queiroz, and L. Vinhas. 2016. Spatiotemporal data representation in R. Pages 178–191 GeoInfo.

Saunders, M. I., M. Bode, S. Atkinson, C. J. Klein, A. Metaxas, J. Beher, M. Beger, M. Mills, R. Giakoumi, and V. Tulloch. 2017. Simple rules can guide whether land-or ocean-based conservation will best benefit marine ecosystems. PLoS biology 15:e2001886.

Saunders, M. I., C. Doropoulos, E. Bayraktarov, R. C. Babcock, D. Gorman, A. M. Eger, M. L. Vozzo, C. L. Gillies, M. A. Vanderklift, A. D. L. Steven, R. H. Bustamante, and B. R. Silliman. 2020. Bright Spots in Coastal Marine Ecosystem Restoration. Current Biology 30:R1500–R1510.

Schiller, A., G. B. Brassington, P. Oke, M. Cahill, P. Divakaran, M. Entel, J. Freeman, D. Griffin, M. Herzfeld, R. Hoeke, X. Huang, E. Jones, E. King, B. Parker, T. Pitman, U. Rosebrock, J. Sweeney, A. Taylor, M. Thatcher, R. Woodham, and A. Zhong. 2020. Bluelink ocean forecasting Australia: 15 years of operational ocean service delivery with societal, economic and environmental benefits. Journal of Operational Oceanography 13:1–18.

Schlaefer, J. A., E. Wolanski, J. Lambrechts, and M. J. Kingsford. 2018. Wind Conditions on the Great Barrier Reef Influenced the Recruitment of Snapper (Lutjanus carponotatus). Frontiers in Marine Science 5.

Shedrawi, G., J. L. Falter, K. J. Friedman, R. J. Lowe, M. S. Pratchett, C. J. Simpson, C. W. Speed, S. K. Wilson, and Z. Zhang. 2017. Localised hydrodynamics influence vulnerability of coral communities to environmental disturbances. Coral Reefs 36:861–872.

Smith, K. A., J. L. Whitney, M. A. McManus, J. Lecky, A. Copeland, D. R. Kobayashi, and J. M. Gove. 2021. Physical mechanisms driving biological accumulation in surface lines on coastal Hawaiian waters. Continental Shelf Research 230:104558.

Sous, D., S. Maticka, S. Meulé, and F. Bouchette. 2022. Bottom Drag Coefficient on a Shallow Barrier Reef. Geophysical Research Letters 49:e2021GL097628.

Sponaugle, S., R. K. Cowen, A. Shanks, S. G. Morgan, J. M. Leis, J. Pineda, G. W. Boehlert, M. J. Kingsford, K. C. Lindeman, and C. Grimes. 2002. Predicting self-recruitment in marine populations: biophysical correlates and mechanisms. Bulletin of Marine Science 70:341–375.

Sudo, K., T. A. L. Quiros, A. Prathep, C. V. Luong, H.-J. Lin, J. S. Bujang, J. L. S. Ooi, M. D. Fortes, M. H. Zakaria, and S. M. Yaakub. 2021. Distribution, temporal change, and conservation status of tropical seagrass beds in Southeast Asia: 2000–2020. Frontiers in Marine Science 8:637722.

Sutherland, G., N. Soontiens, F. Davidson, G. C. Smith, N. Bernier, H. Blanken, D. Schillinger, G. Marcotte, J. Röhrs, K.-F. Dagestad, K. H. Christensen, and Ø. Breivik. 2020. Evaluating the Leeway Coefficient of Ocean Drifters Using Operational Marine Environmental Prediction Systems. Journal of Atmospheric and Oceanic Technology 37:1943–1954.

Swearer, S. E., E. A. Treml, and J. S. Shima. 2019. A Review of Biophysical Models of Marine Larval Dispersal. Pages 325–356 in S. J. Hawkins, A. L. Allcock, A. E. Bates, L. B. Firth, I. P. Smith, S. E. Swearer, and P. A. Todd, editors. Oceanography and Marine Biology. First edition. CRC Press.

Takeyasu, K., Y. Uchiyama, and S. Mitarai. 2023. Quantifying connectivity between mesophotic and shallow coral larvae in Okinawa Island, Japan: a quadruple nested high-resolution modeling study. Frontiers in Marine Science 10:1174940.

Thompson, D. M., J. Kleypas, F. Castruccio, E. N. Curchitser, M. L. Pinsky, B. Jönsson, and J. R. Watson. 2018. Variability in oceanographic barriers to coral larval dispersal: Do currents shape biodiversity? Progress in Oceanography 165:110–122.

Van der Mheen, M., C. Pattiaratchi, S. Cosoli, and M. Wandres. 2020. Depth-dependent correction for wind-driven drift current in particle tracking applications. Frontiers in Marine Science 7:305.

Vanderklift, M. A., C. Doropoulos, D. Gorman, I. Leal, A. J. Minne, J. Statton, A. D. Steven, and T. Wernberg. 2020. Using propagules to restore coastal marine ecosystems. Frontiers in Marine Science:724.

Weeks, R. 2017. Incorporating seascape connectivity in conservation prioritisation. PLOS ONE 12:e0182396.

Wickham, H., and M. H. Wickham. 2017. Package tidyverse. Easily install and load the ‘Tidyverse.

Willis, B. L., and J. K. Oliver. 1990. Direct tracking of coral larvae: Implications for dispersal studies of planktonic larvae in topographically complex environments. Ophelia 32:145–162.

Wolanski, E., T. Asaeda, A. Tanaka, and E. Deleersnijder. 1996. Three-dimensional island wakes in the field, laboratory experiments and numerical models. Continental Shelf Research 16:1437–1452.

Wolanski, E., D. Burrage, and B. King. 1989. Trapping and dispersion of coral eggs around Bowden Reef, Great Barrier Reef, following mass coral spawning. Continental Shelf Research 9:479–496.

Wolanski, E., and W. M. Hamner. 1988. Topographically controlled fronts in the ocean and their biological influence. Science 241:177–181.

Wolanski, E., J. Imberger, and M. L. Heron. 1984. Island wakes in shallow coastal waters. Journal of Geophysical Research: Oceans 89:10553–10569.

Wolanski, E., and B. King. 1990. Flushing of Bowden Reef lagoon, Great Barrier Reef. Estuarine, Coastal and Shelf Science 31:789–804.

Wolanski, E., M. Kingsford, J. Lambrechts, and G. Marmorino. 2024. The Physical Oceanography of the Great Barrier Reef:: A Review. Oceanographic Processes of Coral Reefs:9–34.

Wood, S., I. B. Baums, C. B. Paris, A. Ridgwell, W. S. Kessler, and E. J. Hendy. 2016. El Niño and coral larval dispersal across the eastern Pacific marine barrier. Nature Communications 7:12571.

Zhi, H., J. Siwabessy, S. L. Nichol, and B. P. Brooke. 2014. Predictive mapping of seabed substrata using high-resolution multibeam sonar data: A case study from a shelf with complex geomorphology. Marine Geology 357:37–52.

