## Supplementary Information for "Going with the flow: leveraging reef-scale hydrodynamics for upscaling larval-based restoration"

^2^: CSIRO Environment, St Lucia, Queensland, Australia

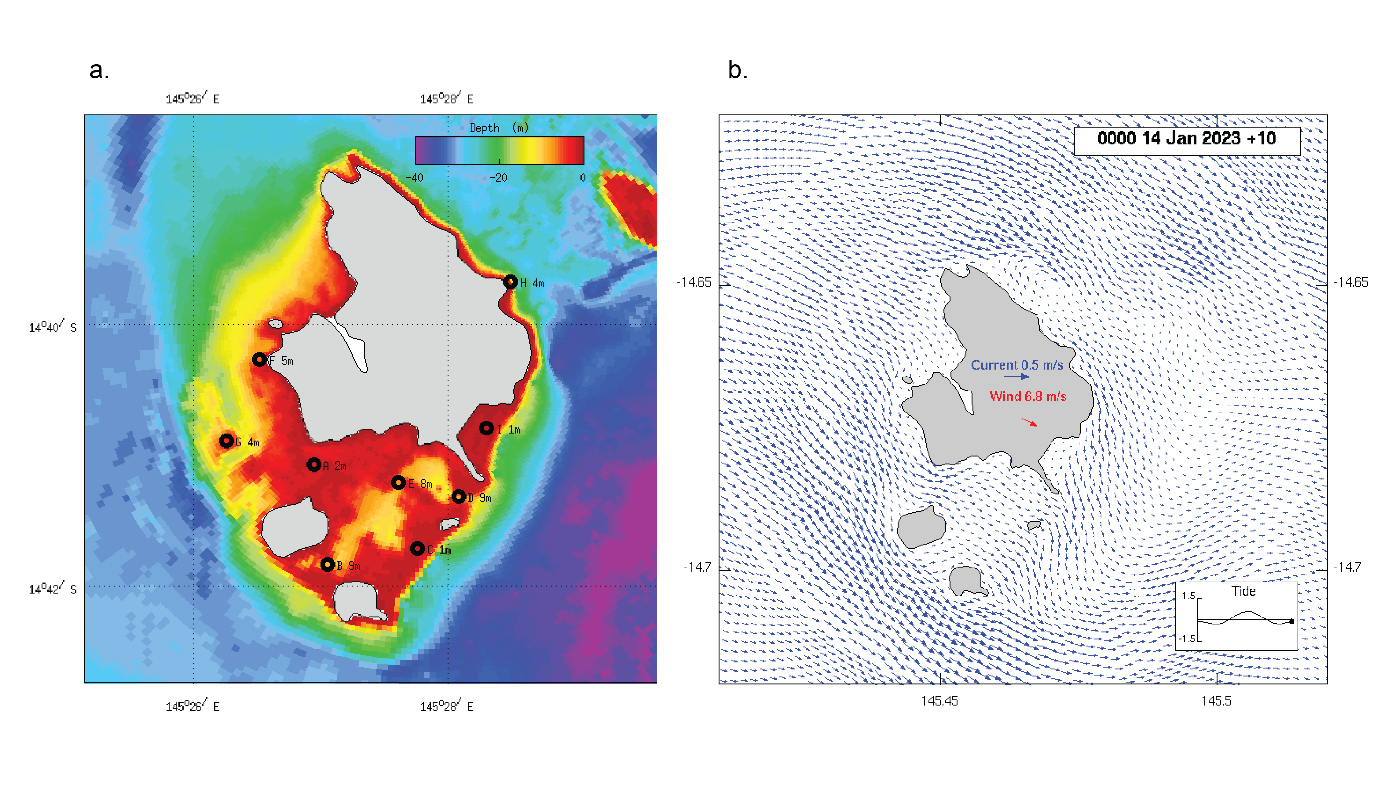

Figure S1: Map of study location showing the fine-resolution bathymetry of Lizard Island reefs (a) and a snapshot in time of surface current conditions around Lizard Island and model boundaries (b)

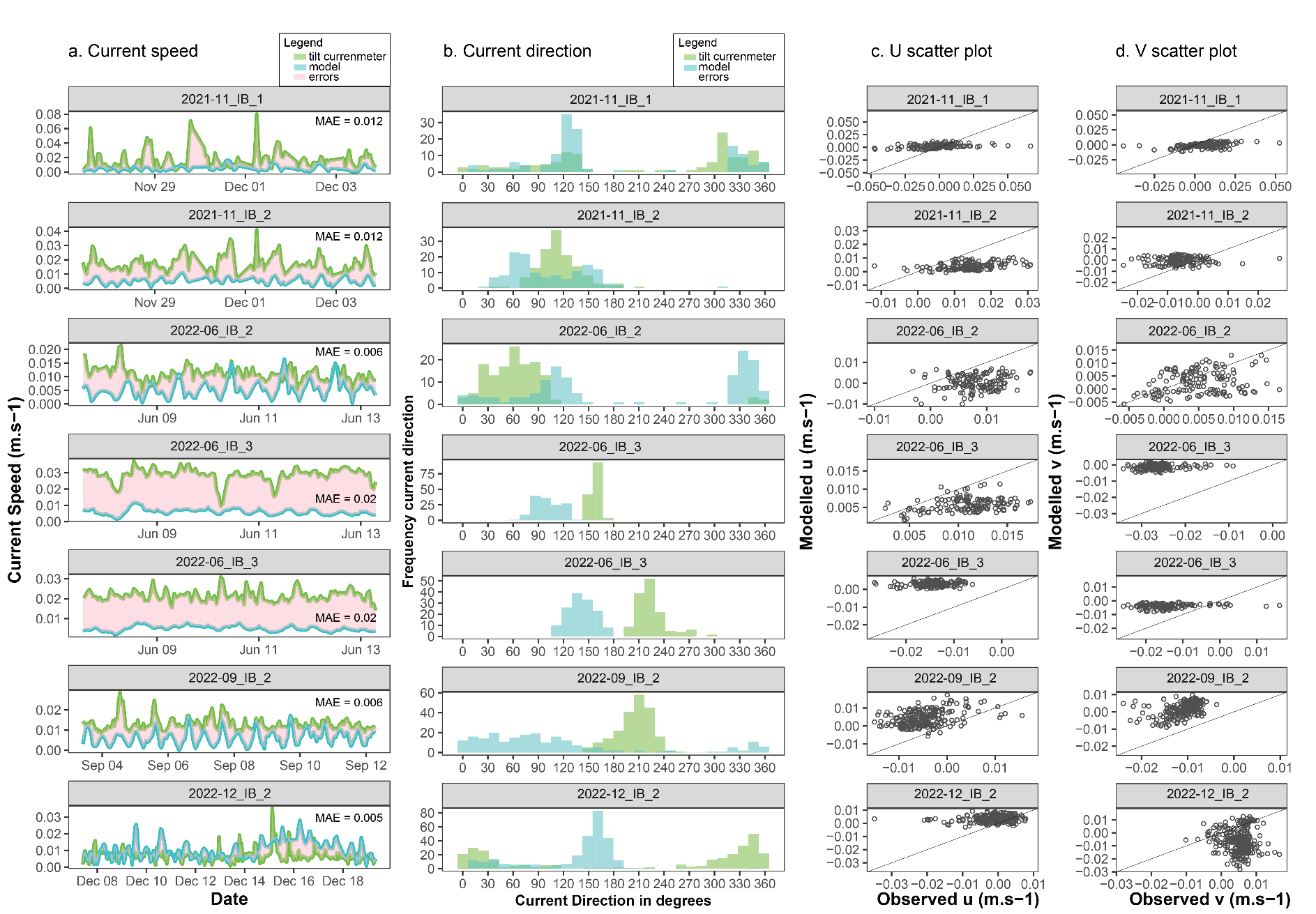

Figure S2: Modelled and observed current speed (a) and direction (b) and scatter plots for u and v components (c,d) for reef locations in the inner bay habitat. All relationships are non significant

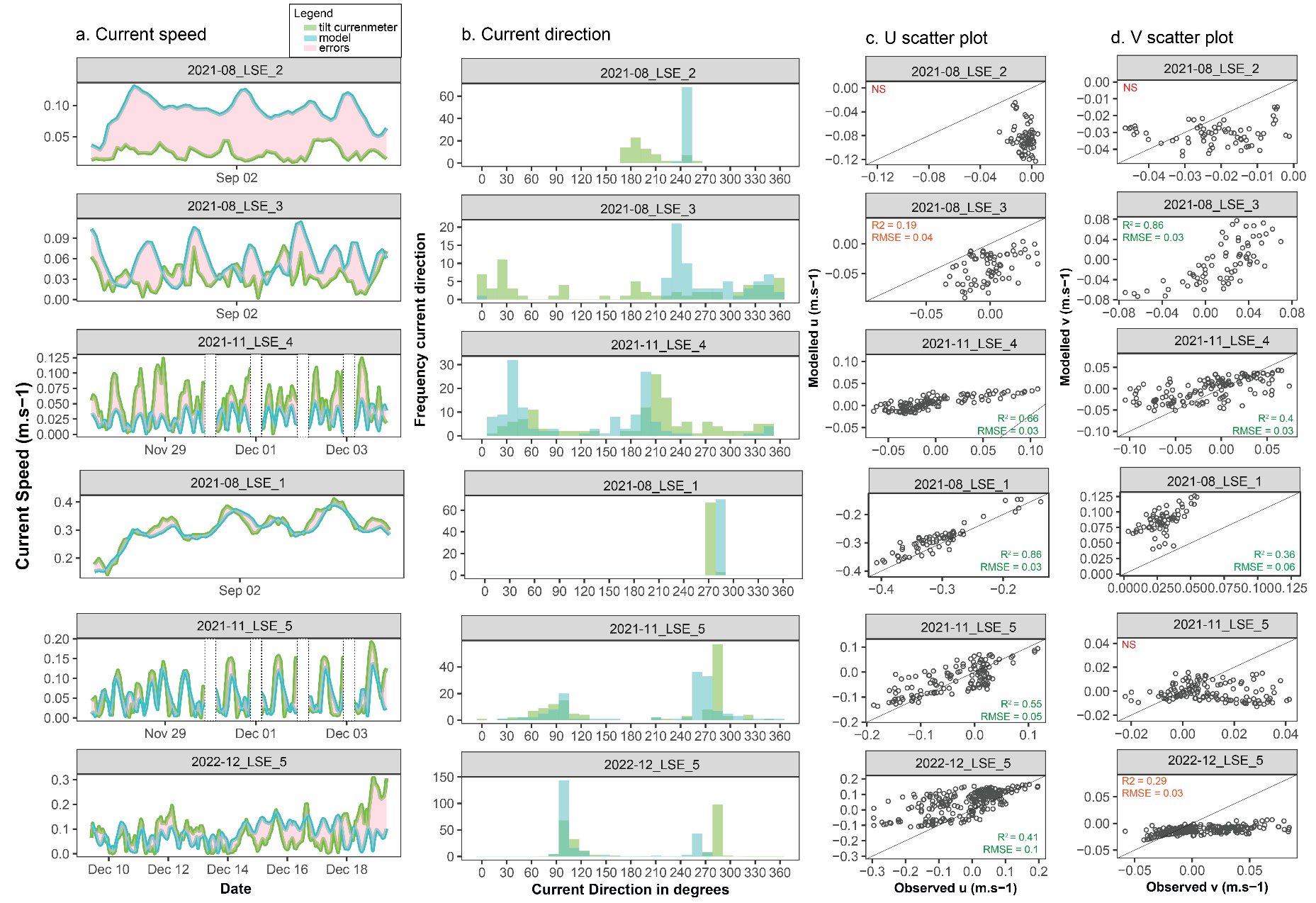

Figure S3: Modelled and observed current speed (a) and direction (b) and scatter plots for u and v components (c,d) for reef locations in the South East Lagoon habitat.

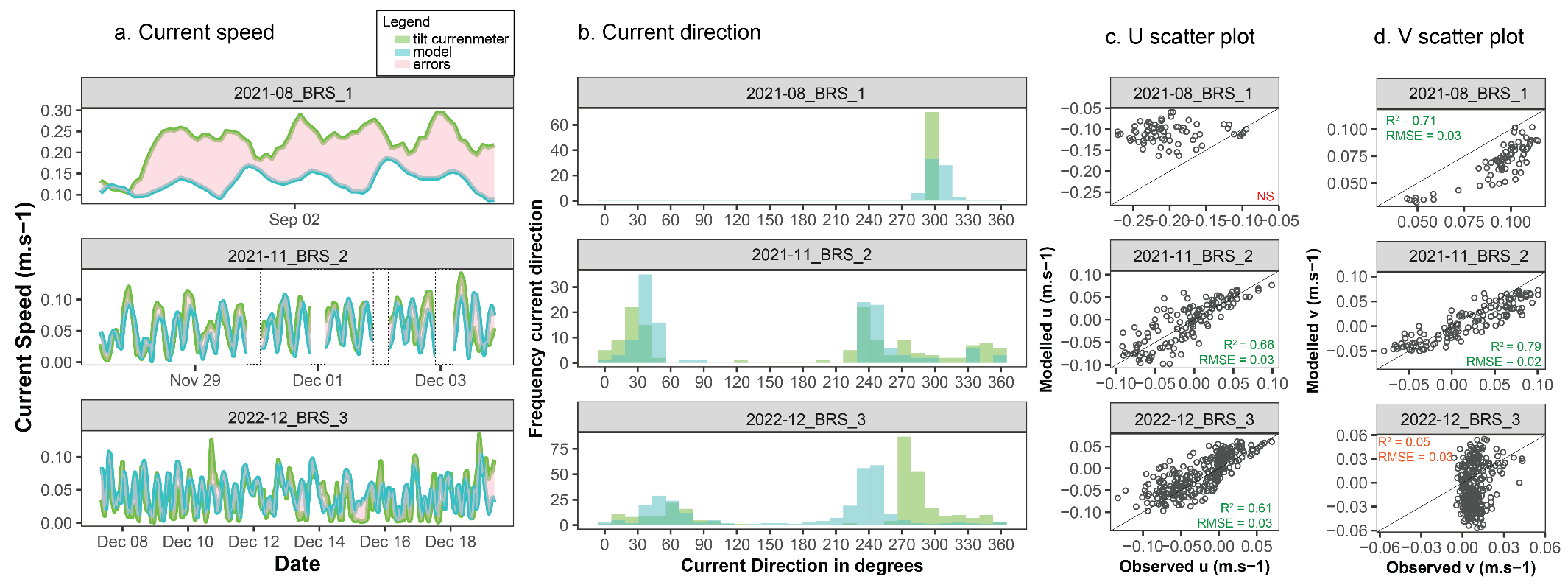

Figure S4: Modelled and observed current speed (a) and direction (b) and scatter plots for u and v components (c,d) for reef locations in the back reef slope habitat.

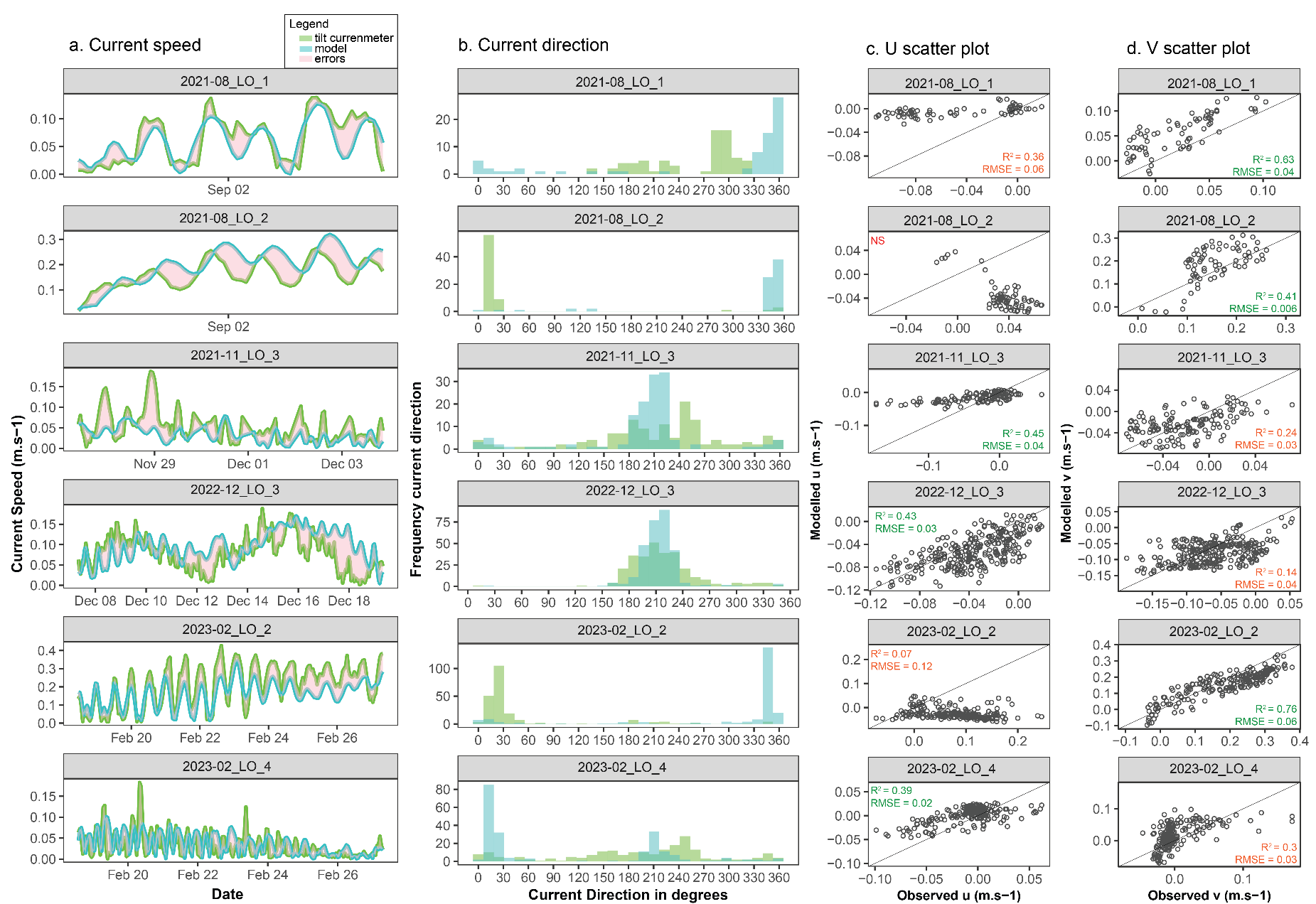

Figure S5: Modelled and observed current speed (a) and direction (b) and scatter plots for u and v components (c,d) for reef locations in the open lagoon habitat.

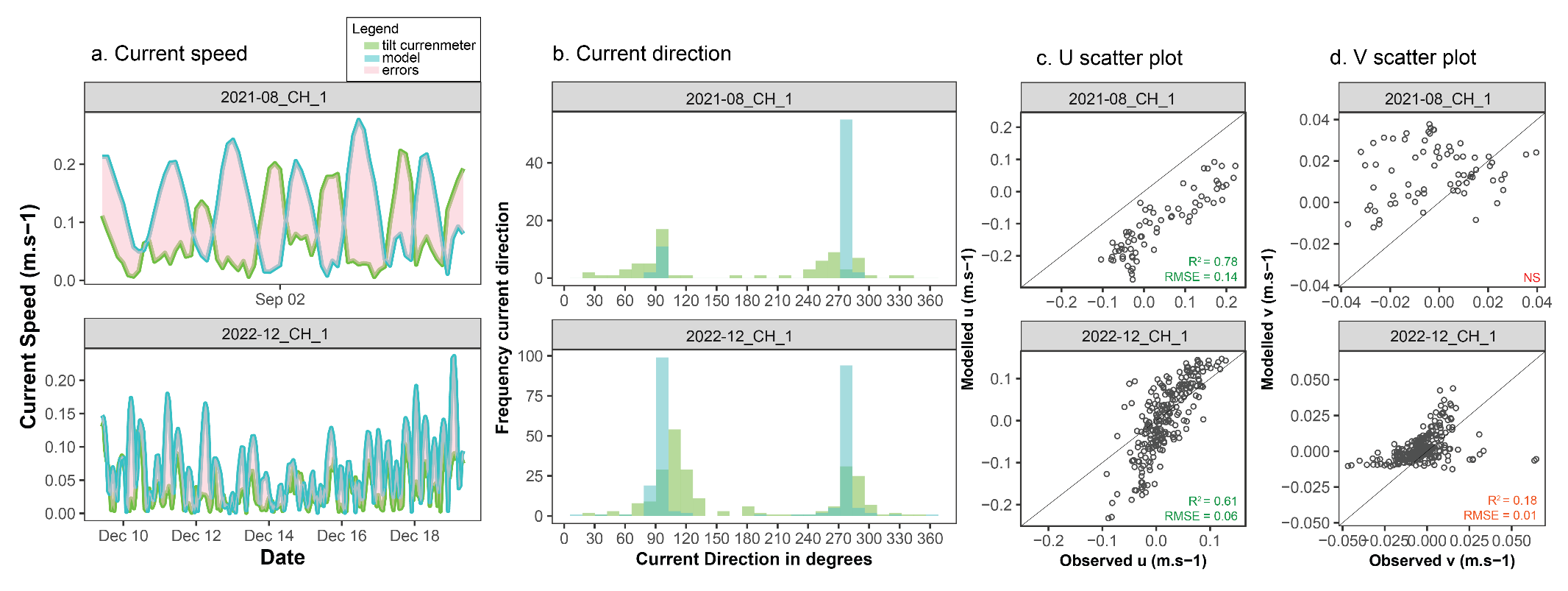

Figure S6: Modelled and observed current speed (a) and direction (b) and scatter plots for u and v components (c,d) for reef locations in the channel habitat.

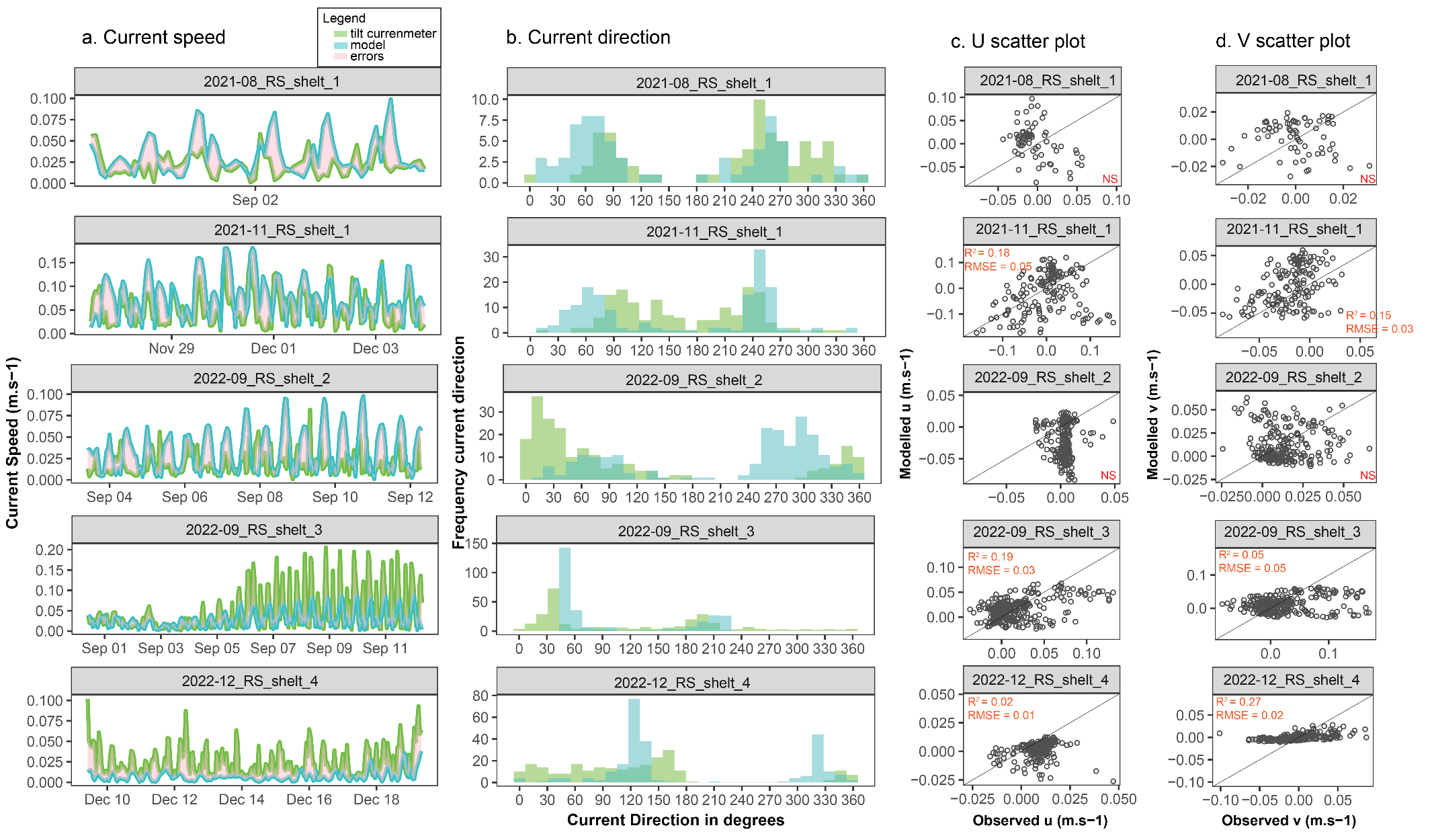

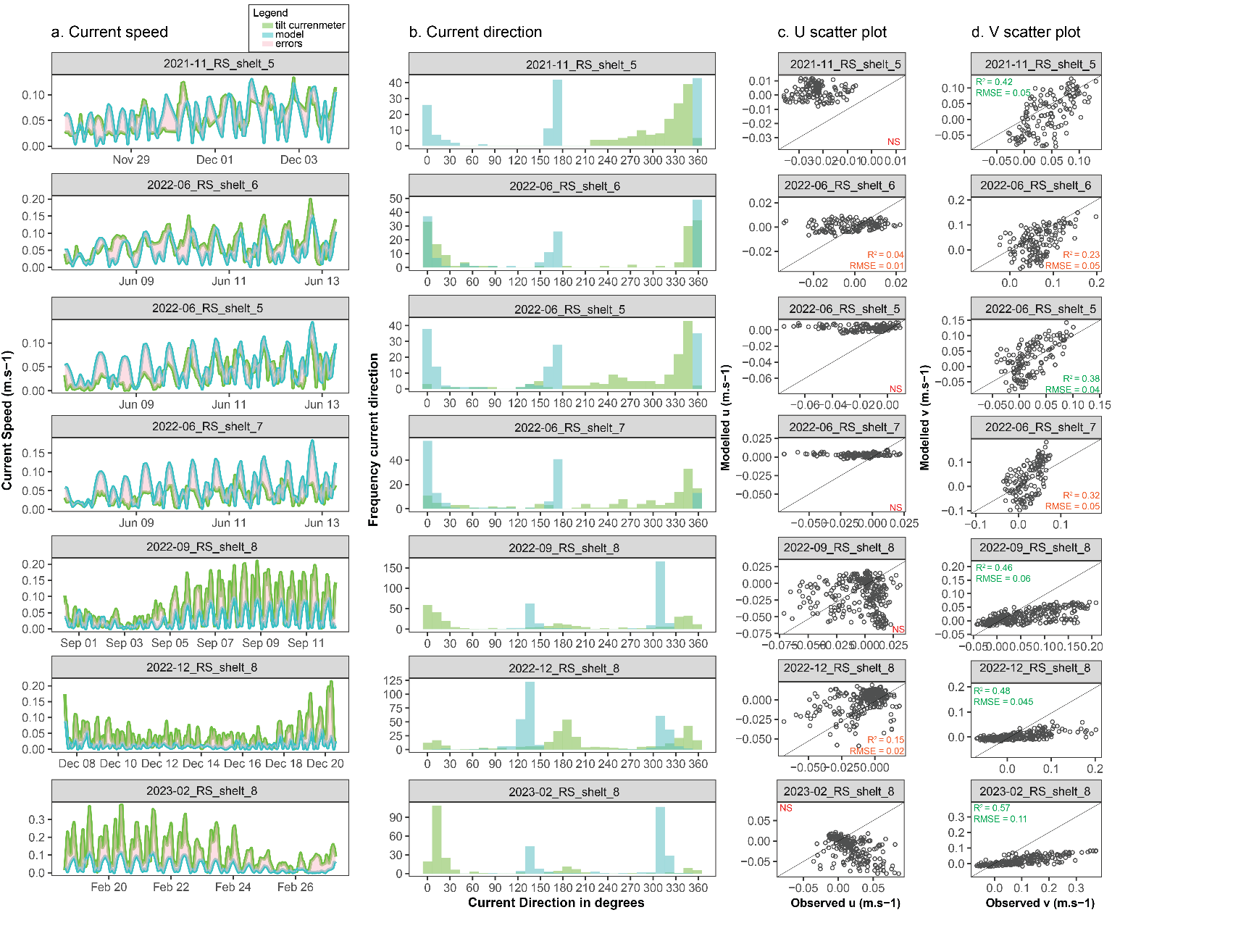

Figure S7: Modelled and observed current speed (a) and direction (b) and scatter plots for u and v components (c,d) for reef locations in the sheltered reef slope habitat.

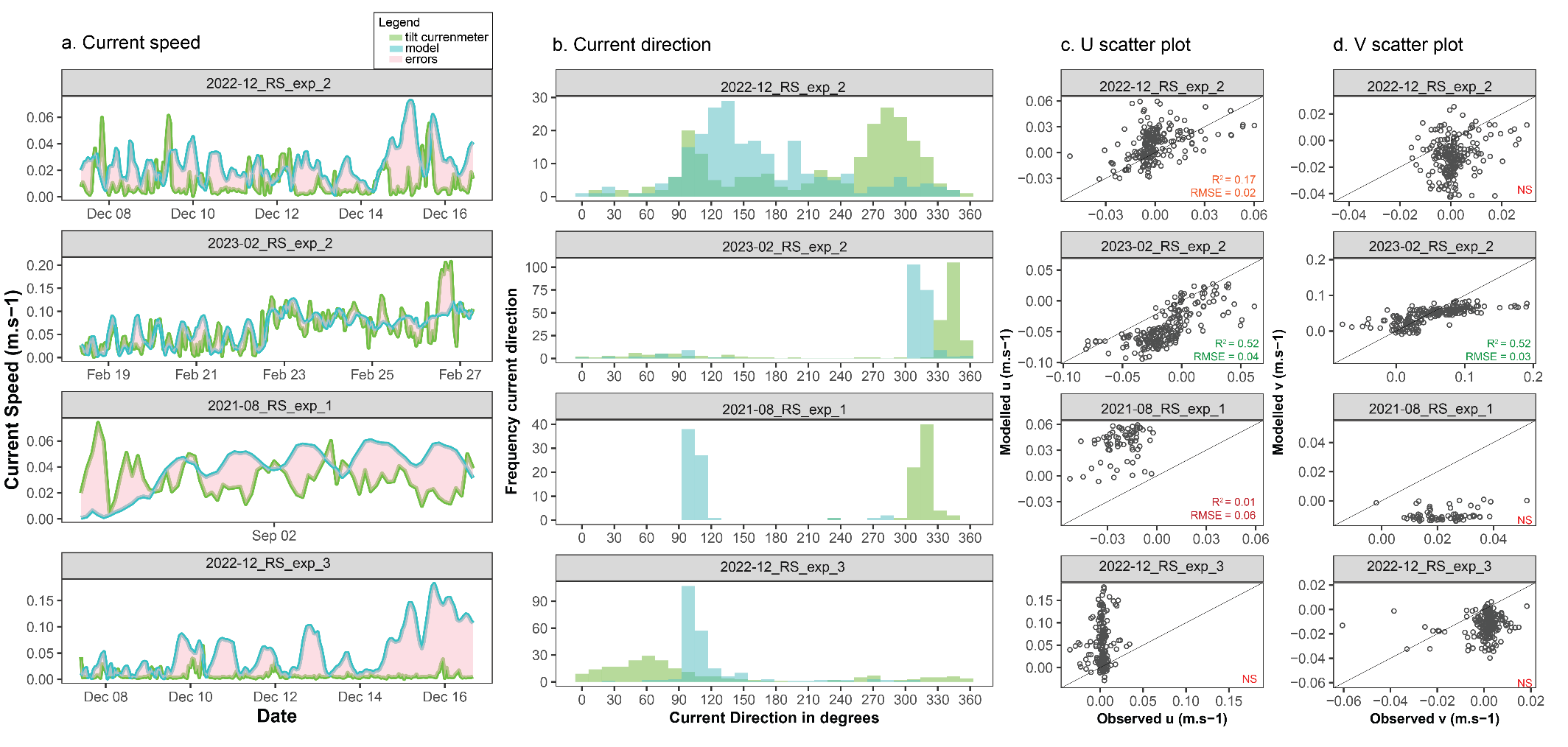

Figure S8: Modelled and observed current speed (a) and direction (b) and scatter plots for u and v components (c,d) for reef locations in the exposed reef slope habitat.

Table S1: Complementary results to Table 1 from the validation analysis below 2 meters depth, comparing modelled current data with tiltmeter current data are several reefs during the study timeframe. Coloured cells highlight the strength of the linear relationship (R2) between modelled and observed velocity vectors; mean errors, RMSE and p-value are also presented. The fit category summarizes the overall representation of local current conditions by the model after detailed visual interpretation (Fig SXX) and expert knowledge explanations.

| Reef Location ID | n | mean_u_err | RMSE_u | R2_u | p_value_u | mean_v_err | RMSE_v | R2_v | p_value_v | Fit Category |
| --- | --- | --- | --- | --- | --- | --- | --- | --- | --- | --- |
| BRS_2 | 135 | 0.003 | 0.028 | 0.660 | 0.000 | -0.011 | 0.024 | 0.797 | 0.000 | Very Good Fit |
| BRS_3 | 289 | 0.011 | 0.028 | 0.615 | 0.000 | -0.014 | 0.029 | 0.051 | 0.000 | Very Good Fit |
| LO_1 | 70 | 0.044 | 0.057 | 0.361 | 0.000 | 0.034 | 0.041 | 0.632 | 0.000 | Very Good Fit |
| LO_2 | 214 | -0.107 | 0.126 | 0.073 | 0.000 | -0.031 | 0.063 | 0.767 | 0.000 | Very Good Fit |
| LSE_1 | 70 | 0.020 | 0.028 | 0.859 | 0.000 | 0.057 | 0.059 | 0.363 | 0.000 | Very Good Fit |
| LSE_4 | 137 | 0.009 | 0.030 | 0.661 | 0.000 | 0.013 | 0.036 | 0.402 | 0.000 | Very Good Fit |
| LSE_5 | 137 | 0.007 | 0.046 | 0.548 | 0.000 | -0.011 | 0.019 | 0.014 | 0.171 | Very Good Fit |
| RS_exp_2 | 215 | -0.031 | 0.037 | 0.523 | 0.000 | -0.008 | 0.034 | 0.524 | 0.000 | Very Good Fit |
| CH_1 | 70 | -0.141 | 0.148 | 0.782 | 0.000 | 0.017 | 0.026 | 0.038 | 0.107 | Good Fit |
| CH_1 | 239 | -0.011 | 0.059 | 0.611 | 0.000 | 0.005 | 0.014 | 0.178 | 0.000 | Good Fit |
| LO_2 | 70 | -0.075 | 0.084 | 0.590 | 0.000 | 0.033 | 0.067 | 0.419 | 0.000 | Good Fit |
| LO_3 | 137 | 0.015 | 0.040 | 0.460 | 0.000 | -0.004 | 0.027 | 0.249 | 0.000 | Good Fit |
| LO_3 | 289 | -0.015 | 0.028 | 0.434 | 0.000 | -0.016 | 0.046 | 0.196 | 0.000 | Good Fit |
| LO_4 | 210 | 0.010 | 0.024 | 0.394 | 0.000 | 0.013 | 0.036 | 0.303 | 0.000 | Good Fit |
| LSE_3 | 70 | -0.038 | 0.044 | 0.199 | 0.000 | -0.022 | 0.036 | 0.510 | 0.000 | Good Fit |
| LSE_5 | 239 | 0.070 | 0.105 | 0.417 | 0.000 | -0.019 | 0.032 | 0.288 | 0.000 | Good Fit |
| RS_shelt_8 | 287 | -0.012 | 0.034 | 0.003 | 0.389 | -0.041 | 0.064 | 0.469 | 0.000 | Good Fit |
| RS_shelt_8 | 315 | 0.006 | 0.016 | 0.157 | 0.000 | -0.013 | 0.045 | 0.482 | 0.000 | Good Fit |
| RS_shelt_8 | 210 | -0.041 | 0.059 | 0.249 | 0.000 | -0.080 | 0.112 | 0.579 | 0.000 | Good Fit |
| RS_shelt_3 | 286 | -0.002 | 0.030 | 0.196 | 0.000 | -0.010 | 0.051 | 0.049 | 0.000 | Good Fit |
| RS_shelt_5 | 141 | 0.020 | 0.027 | 0.001 | 0.693 | 0.004 | 0.041 | 0.380 | 0.000 | Good Fit_wind |
| RS_shelt_6 | 141 | 0.003 | 0.011 | 0.045 | 0.011 | -0.034 | 0.058 | 0.231 | 0.000 | Good Fit_wind |
| RS_shelt_7 | 141 | 0.014 | 0.022 | 0.000 | 0.994 | 0.006 | 0.055 | 0.326 | 0.000 | Good Fit_wind |
| BRS_1 | 70 | 0.092 | 0.103 | 0.004 | 0.620 | -0.024 | 0.026 | 0.714 | 0.000 | Poor Fit |
| RS_exp_1 | 70 | 0.059 | 0.061 | 0.106 | 0.006 | -0.032 | 0.033 | 0.042 | 0.089 | Poor Fit |
| RS_exp_2 | 225 | 0.010 | 0.020 | 0.174 | 0.000 | -0.014 | 0.019 | 0.048 | 0.001 | Poor Fit |
| IB_1 | 136 | 0.003 | 0.016 | 0.133 | 0.000 | -0.006 | 0.013 | 0.229 | 0.000 | No Fit |
| IB_2 | 136 | -0.010 | 0.012 | 0.134 | 0.000 | 0.006 | 0.009 | 0.001 | 0.767 | No Fit |
| IB_2 | 139 | -0.008 | 0.009 | 0.035 | 0.027 | -0.002 | 0.006 | 0.061 | 0.003 | No Fit |
| IB_2 | 214 | 0.009 | 0.010 | 0.147 | 0.000 | 0.013 | 0.013 | 0.208 | 0.000 | No Fit |
| IB_2 | 287 | 0.005 | 0.007 | 0.028 | 0.005 | -0.012 | 0.015 | 0.006 | 0.197 | No Fit |
| IB_3 | 139 | -0.005 | 0.006 | 0.088 | 0.000 | 0.025 | 0.025 | 0.025 | 0.064 | No Fit |
| IB_4 | 139 | 0.017 | 0.018 | 0.018 | 0.120 | 0.010 | 0.012 | 0.031 | 0.039 | No Fit |
| LSE_2 | 70 | -0.080 | 0.084 | 0.050 | 0.063 | -0.009 | 0.015 | 0.014 | 0.336 | No Fit |
| RS_exp_3 | 223 | 0.046 | 0.065 | 0.011 | 0.125 | -0.013 | 0.017 | 0.018 | 0.044 | No Fit |
| RS_shelt_1 | 70 | 0.007 | 0.051 | 0.181 | 0.000 | 0.002 | 0.018 | 0.008 | 0.458 | No Fit |
| RS_shelt_1 | 158 | -0.015 | 0.087 | 0.056 | 0.003 | 0.015 | 0.036 | 0.152 | 0.000 | No Fit |
| RS_shelt_2 | 214 | -0.025 | 0.039 | 0.003 | 0.443 | -0.002 | 0.024 | 0.001 | 0.578 | No Fit |
| RS_shelt_4 | 239 | -0.007 | 0.012 | 0.024 | 0.016 | 0.001 | 0.025 | 0.272 | 0.000 | No Fit |
| RS_shelt_5 | 136 | 0.025 | 0.026 | 0.007 | 0.334 | -0.026 | 0.052 | 0.424 | 0.000 | No Fit |

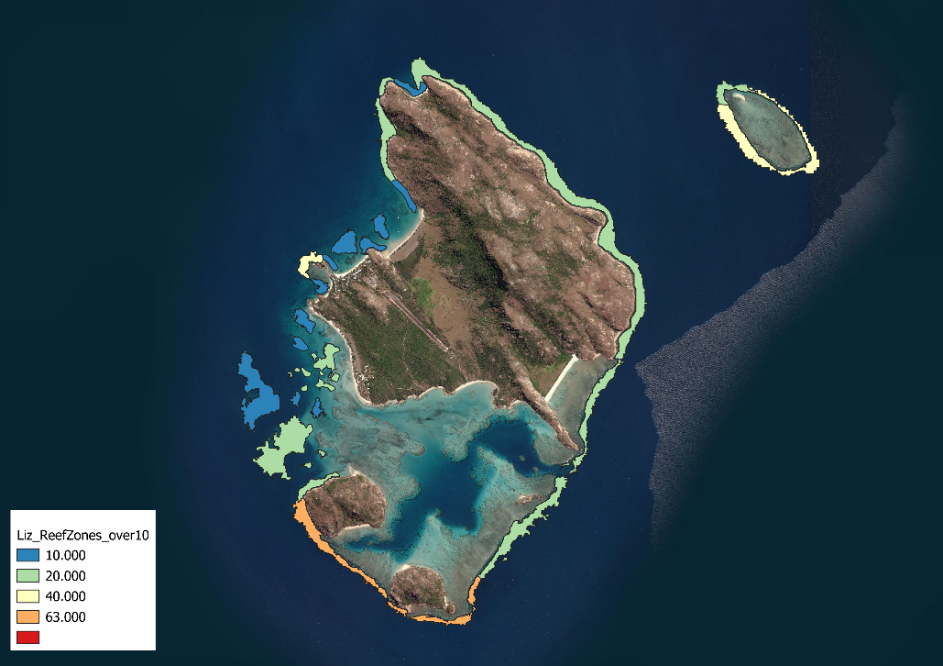

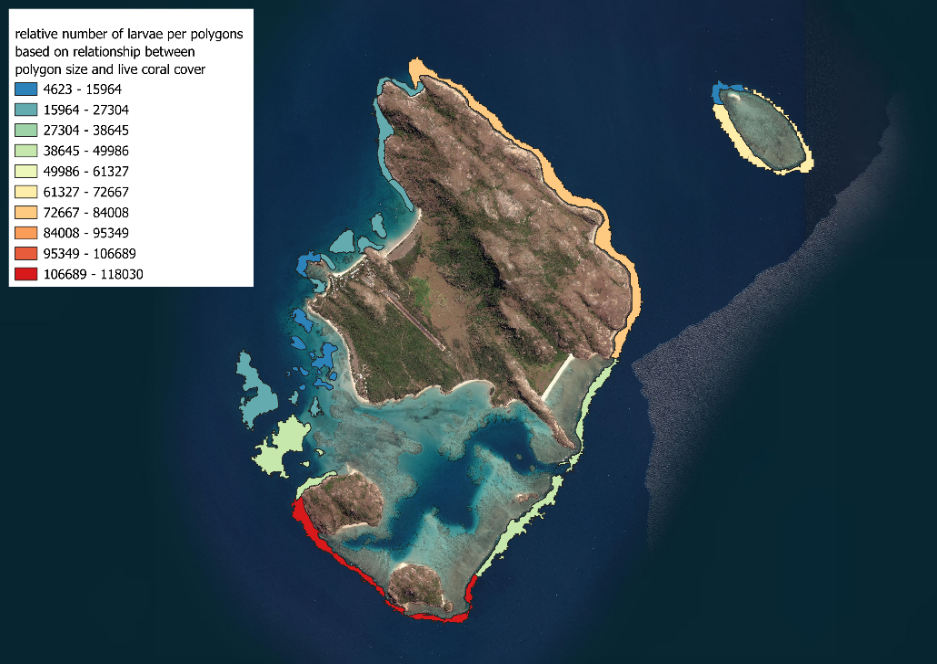

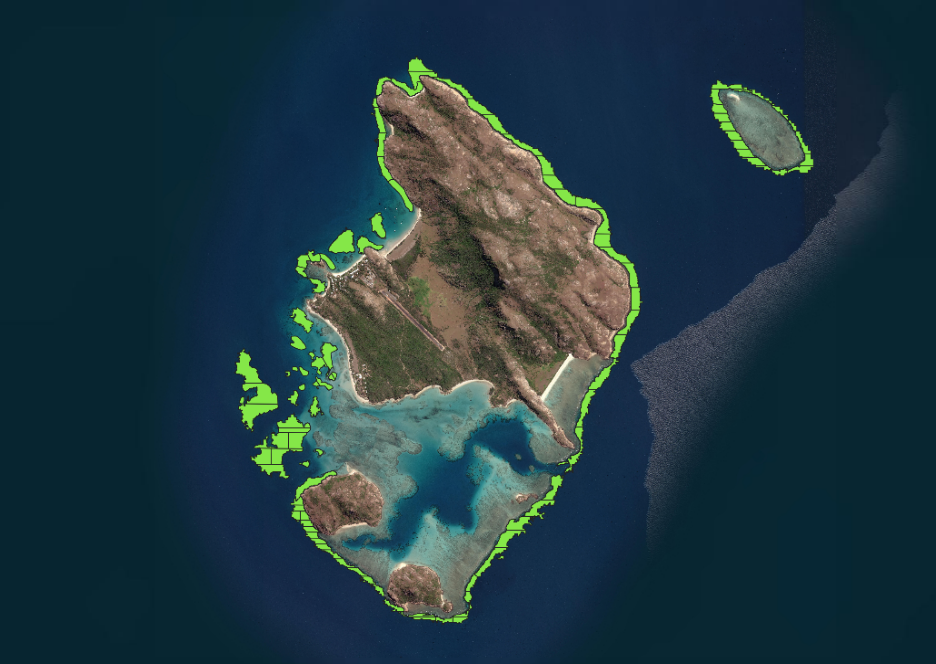

Figure S9: GIS workflow showing the relation between live coral cover (top), relative number of larvae after Roff (2023) and polygons divided into equal number of larvae for modelling exercise (bottom).

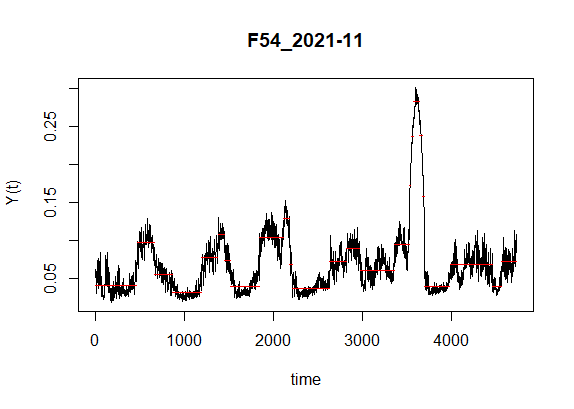

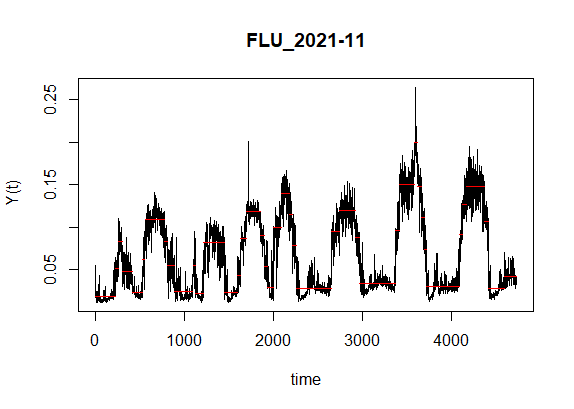

Figure S10: Examples of time series of tiltcurrentmeters at two different sites and the timeframes highlighted in red showing significant change in current speed. The changepoint analysis was used to detect timeframes of slack current conditions

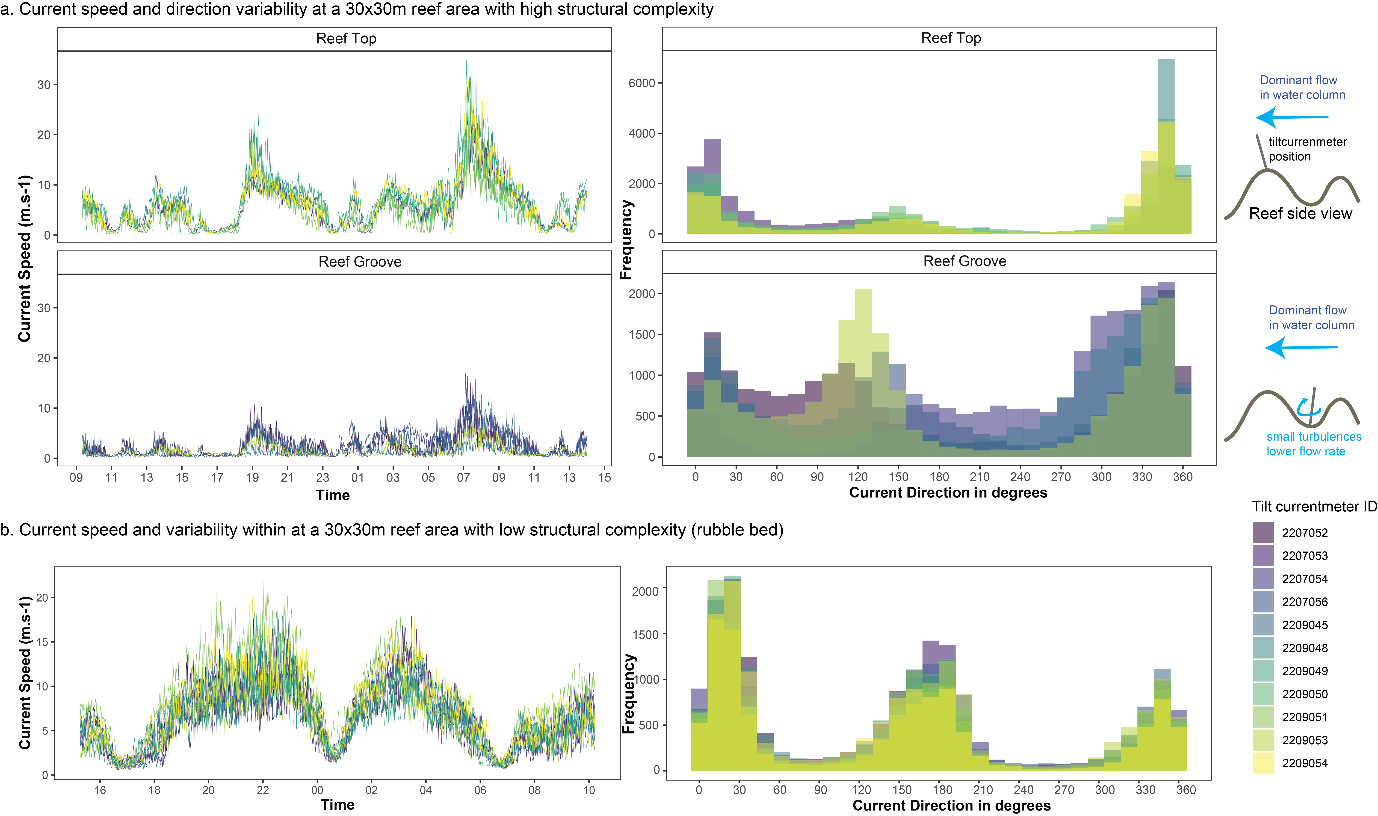

Figure S11. 24-hour current speed trends and frequency distribution of current direction collected by 11 tilt currentmeters spread within a model grid cell (ie. 30 m x 30 m reef area) at a structurally complex reef (a) and a rubble bed with low structural complexity (b). (a) shows the significant influence of complexity in adding variability in observed current speed and direction on a reef. Note different scales on y-axes when comparing plots.

a. Particle’s residence time at the restoration site as a function of the timing of delivery

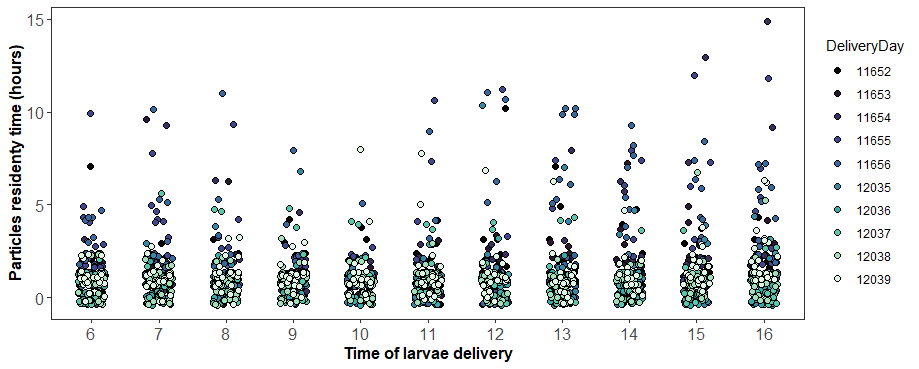

b. Particle’s residence time at the 25 restoration sites

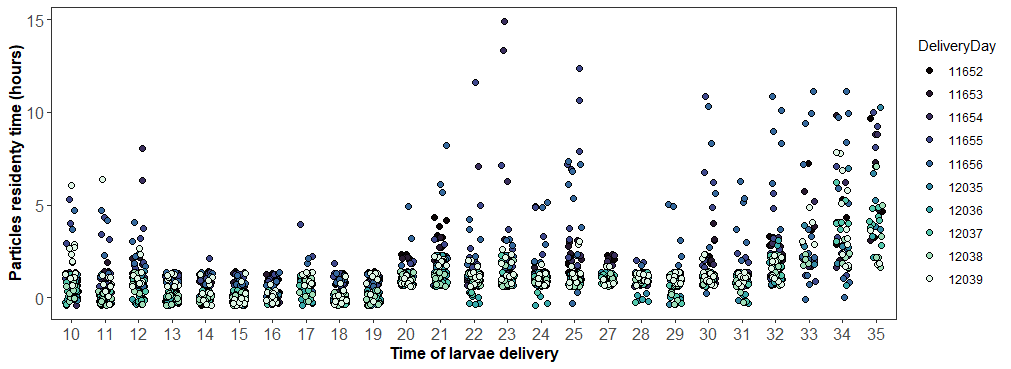

Figure S12: plot showing the particles' residency time in hours for each delivery time and each delivery sites (#10 to #35)

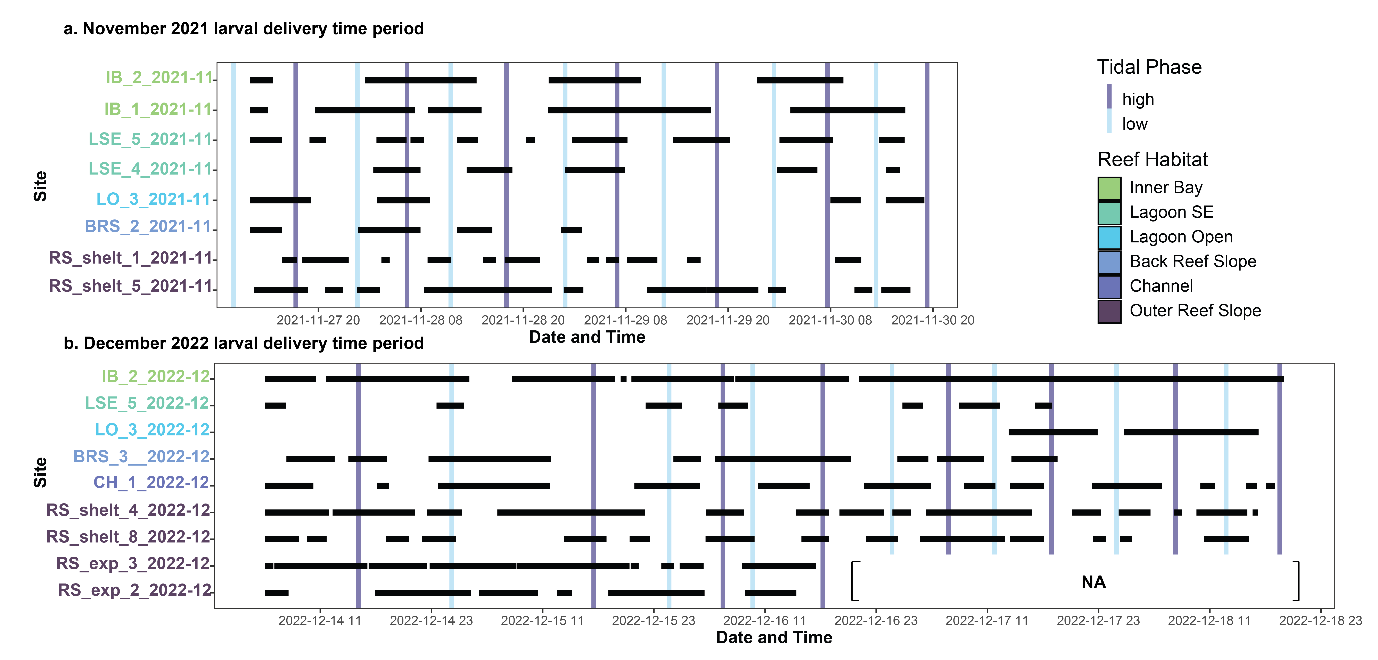

Figure S13: Timeframes of slack current conditions on reefs in different habitat during larval delivery in 2021 (a) and 2022 (b). Timing of high and low tidal phase are represented with vertical purple and blue segments, respectively. Slack periods are represented with horizontal black segments. The end of the time series at RS_exp 2 and 3 is not available due to instrument retrieval before bad weather.

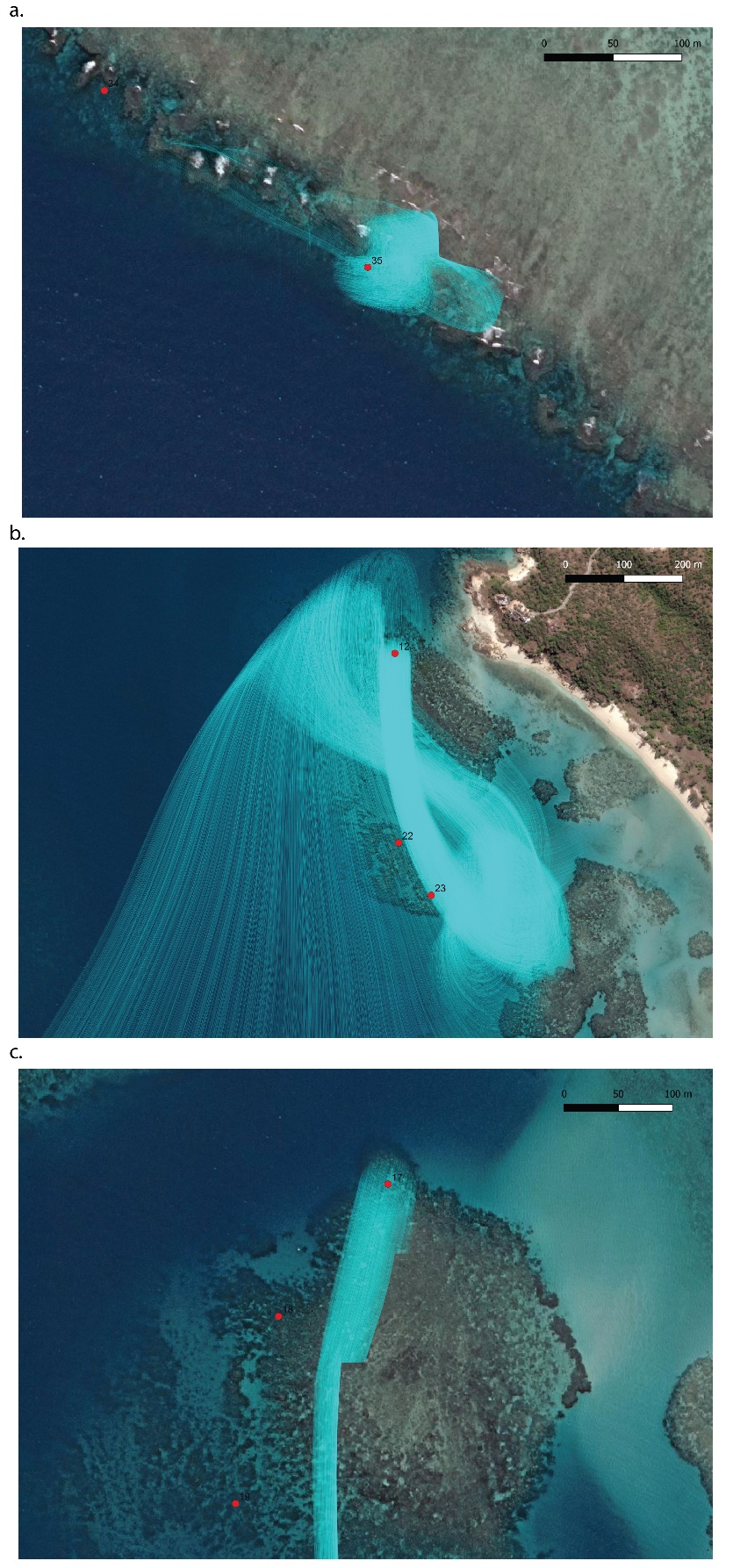

Figure S14: Examples of trajectories of the 5000 particles through time for some scenarios showing high PRT at site 35 (a), site 12 (b). Red dots shows the center of the 25m radius delivery polygon

**Appendix 1:** Larval delivery phase: predicting the residency of particles over damaged reefs

In certain simulation scenarios, a significant proportion of particles remained stationary throughout the 24-hour simulation period, indicating 'stuck' particles.

If more than 30% of particles in a simulation exhibited stationary behaviour that lasted over 1 hour, the results from those simulations were excluded from subsequent analyses.

If less than 30% of particles displayed stationary behaviour in a simulation, these stationary particles were filtered out from the dispersal output. The remaining data, representing mobile particles, were retained for further analysis. This resulted in a dataset with spatial and temporal information for 3,500 particles, as opposed to the original 5000 particles

Dispersal output .json files examples, codes, and data output examples for larval delivery phase are all accessible on Marine Gouezo’s mgouezo github public repository called ‘LarvalDeliverySim’

<https://github.com/mgouezo/LarvalDeliverySim>
